# Inflammatory signaling differentially changes chromatin accessibility and gene expression of the PD- associated kinase LRRK2 between human and mice

**DOI:** 10.64898/2026.01.11.698894

**Authors:** Alexandra Beilina, Jae-Hyeon Park, Natalie Landeck, Ruth Chia, Jinhui Ding, Xylena Reed, Hannah Bailey, Changyoun Kim, Alice Kaganovich, Dominic J. Acri, Janet Brooks, Alexandra Mann, Kareem A. Zaghloul, Elise Marsan, J. Raphael Gibbs, Alex DeCasien, Mark R. Cookson

**Author notes:** These authors contributed equally (Alexandra Beilina, Jae-Hyeon Park, Natalie Landeck).

## Abstract

The genomic locus that encodes the Leucine-rich repeat kinase 2 (LRRK2) gene is highly pleiotropic, being associated with both Parkinson’s disease (PD) and Crohn’s disease (CD). Coding variants associated with risk of PD and CD act as gain of function kinase mutations increasing phosphorylation of RAB substrates, and non-coding variants in the promoter region of LRRK2 increase expression of the gene, notably in immune cells. If regulation of LRRK2 expression is a causal contributor to age-related diseases, it is important to understand the mechanism(s) by which LRRK2 is regulated, particularly in the context of inflammation. Here, we show that interferon-ɣ exposure induces robust LRRK2 activation in human iPSC-derived microglia through signaling of the Janus-activated Kinase complex to phosphorylate STAT1, which then binds to the LRRK2 promoter and is associated with remodeling of chromatin structure in this genomic locus. Additional regulatory mechanisms include the stress-induced transcription factor and long non-coding RNA encoded at the same locus, resulting in increased LRRK2 mRNA levels. We also show evidence of the same effect in acutely cultured human brain slices. While we were unable to demonstrate any induction of Lrrk2 mRNA in the mouse brain, the introduction of a human bacterial artificial chromosome transgene into the mouse genome recapitulated sensitivity to interferon-ɣ in microglia. A comparative genomic analysis across mammals suggests that these species differences are driven by regulatory regions upstream of LRRK2 that are specific to anthropoid primates. These results demonstrate that there are differences between species in how genes associated with human diseases are regulated and provide important information that should be incorporated in disease modeling.

## Introduction

Complex and common diseases are proposed to be highly polygenic, that is with risk alleles distributed across many loci (1). Furthermore, it is postulated that this polygenic risk interacts with demographic and environmental factors to establish diseases, especially those of middle and later life (2). In turn, aging is associated with multiple molecular changes, not limited to but including chronic inflammation (3). These considerations have led to a proposed framework for thinking about age-related neurodegenerative conditions where chronic inflammatory changes are an important disease hallmark (4).

Leucine-rich repeat kinase 2 (*LRRK2*) has emerged as a key player in both neurodegenerative and inflammatory conditions. Mutations in *LRRK2* are a relatively common genetic cause of dominantly inherited Parkinson’s disease (PD) (5–8) and non-coding risk variants have been associated with sporadic PD by genome wide association studies (9). LRRK2 has also independently been nominated as a risk factor for Crohn’s disease (CD) (10,11), a type of inflammatory bowel disease (IBD), which is characterized by chronic inflammation of the gastrointestinal tract (12). These genetic associations suggest that LRRK2 has functional roles in immune cells and implies that PD may also be a disease of chronic inflammation (13).

LRRK2 is expressed in both peripheral immune cell subsets, including monocytes and macrophages (14) that can be recruited to the brain during inflammatory events (15,16). Whether LRRK2 is expressed in parenchymal microglia, critical immune cells in the brain, is less clear. We have previously reported using single-nucleus RNA-seq analyses that microglial expression of LRRK2 is dependent on the genotype of the donor, with PD-associated risk alleles having higher expression than protective alleles (17). Furthermore, we were able to replicate this association in induced pluripotent stem cell (iPSC)-derived microglia (iMicroglia) and showed that genotype-dependent expression propagates to a known cellular function of LRRK2, namely phosphorylation of membrane-associated RAB proteins (18) which occurs after stimuli that induce lysosomal damage (19), including exposure to infectious agents (20, 21).

LRRK2 expression is upregulated in response to inflammatory stimuli, including interferon-ɣ (IFN-ɣ) exposure, in mouse or human monocyte cell lines (22) and in human iMicroglia and dopamine neurons (23). More recently, IL-4 has also been shown to upregulate LRRK2 activity in murine peripheral immune cells (24). This latter observation is particularly intriguing as it has long been thought that IFN-ɣ and IL-4 have opposing roles in immune regulation (25) and might suggest that LRRK2 regulation is more complex than a simple association with pro- or anti- inflammatory conditions.

Collectively, these considerations suggest that a better understanding of the mechanism(s) by which LRRK2 expression is regulated by inflammatory signaling might help identify common pathways between PD and CD. We therefore performed a detailed analysis of how LRRK2 is regulated by IFN-ɣ treatment in iMicroglia. We identify changes in the chromatin structure of the LRRK2 locus and show that IFN-ɣ regulated expression of LRRK2 depends on Janus Activated Kinase - signal transducer and activator of transcription 1 (JAK-STAT1) signaling, supplemented by the transcription factor ATF5 and by long noncoding RNAs at the same locus. We did not see induction of Lrrk2 by Ifn-ɣ in microglia in the mouse brain and find that, across multiple peripheral immune cells, human cells show more robust induction than their murine equivalents. We were able to demonstrate LRRK2 induction by IFN-ɣ in microglia in acutely cultured human brain slices and show that introduction of a bacterial artificial chromosome (BAC) transgene containing human regulatory elements that bind STAT1 is sufficient to infer IFN-ɣ inducibility of LRRK2 in murine microglia. These results suggest that the inflammatory regulated expression of LRRK2 in CNS and peripheral immune cells are likely to be species dependent. In line with this, our comparative genomic analysis across 32 mammalian species detected a specific region of the IFN-γ-LRRK2 regulatory complex that may be specific to anthropoid primates. Differences in LRRK2 regulation and expression that are critical to account for in disease modeling and may partially explain why humans are prone to PD, but other species are not.

## Results

### LRRK2 expression and activity are upregulated by interferon-ɣ in human cells

We first differentiated human iPSC into iMicroglia from the cord-blood derived A18945 female iPSC line using an embryoid body-based protocol (26) as shown schematically in Fig. 1a. Consistent with prior reports (23), we observed a marked increase in LRRK2 protein levels in iPSC-derived microglia upon treatment with 20 ng/mL (equivalent to 200 IU/mL) IFN-ɣ for 24h (Fig. 1b, c). In contrast, the anti-inflammatory agents IL-4 and IL-10 did not affect LRRK2 expression in the same cells (Fig. 1b, c). Levels of pRAB10, a known LRRK2 substrate, were increased by IFN-ɣ but not either of the interleukins (Fig. 1b, d) We found that increases in both LRRK2 protein and the substrate pRAB10 occurred exclusively in response to the group II interferon IFN-ɣ and not group I (IFN-β) or group III (IFN-λ) interferons (Fig. 1e-g). We further examined the specificity of other proinflammatory agents for LRRK2 activation. Both lipopolysaccharide (LPS) and synuclein preformed fibrils (PFFs) significantly elevated LRRK2 and pRAB10 levels, although to a lesser extent than IFN-ɣ (Supplementary Fig. S1a-c). In contrast, stimulating phagocytosis with pHrodo-Zymozan did not affect LRRK2 expression or pRAB10 accumulation (Supplementary Fig. S1a-c).

**Figure 1.**
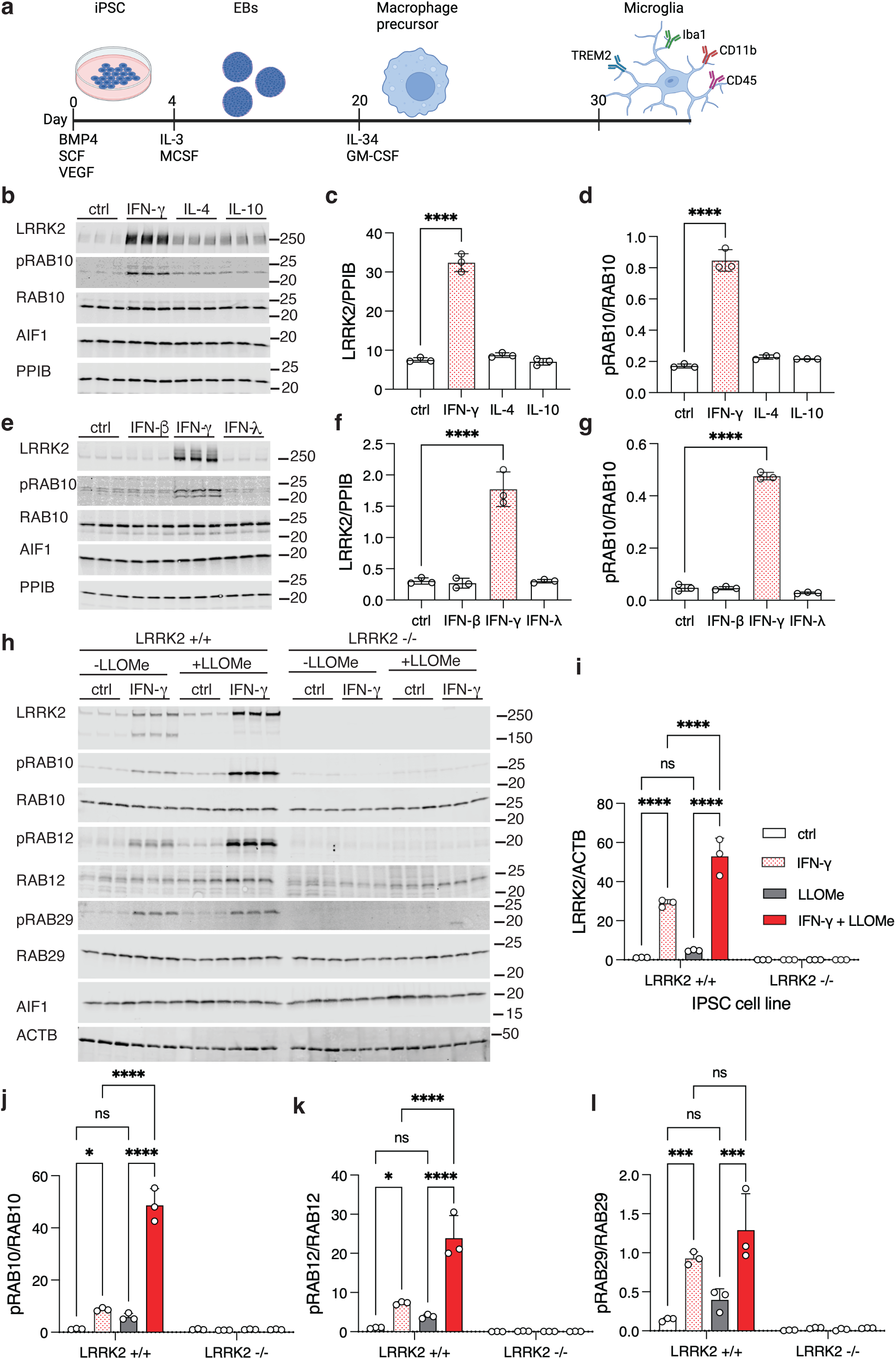
Induction of LRRK2 expression and activity by IFN-ɣ. (a) Schematic showing the induction protocol used to make iPSC-derived microglia (iMicroglia) in this study. (b-d) LRRK2 induction by IFN-ɣ but not anti-inflammatory cytokines in human iMicroglia. (b) iMicroglia were exposed to IFN-ɣ, IL-4 or IL-10 or control media (ctrl) for 24 hours prior to protein extraction and blotting for LRRK2, pRAB10, total RAB10, the microglial marker AIF1 (Iba1) or PPIB (cyclophilin-B) as a protein reference. Quantification of LRRK2 relative to PPIB (c) or pRAB10 relative to total RAB10 (d) shows that LRRK2 protein levels and activity are significantly increased by IFN-ɣ and not other treatments. ***, *p*<0.0001 by one-way ANOVA (F_3,8_=274 for LRRK2, F_3,8_=242 for pRAB10) with Dunnet’s post-hoc test versus control conditions. (e-g). LRRK2 induction by type II interferon. Cells were exposed to IFN-β, IFN-ɣ, or IFN-λ that are representative of type I, type II or type III interferons respectively and blotted for protein markers as in the prior experiment (e) and quantified for LRRK2 (f) and pRAB10 (g). ***, *p*<0.0001 by one-way ANOVA (F_3,8_=77.69 for LRRK2, F_3,8_=1386 for pRAB10) with Dunnet’s post-hoc test versus control conditions. (h-l). LRRK2 expression and induction by combined IFN-ɣ and lysosomal damage. iMicroglia that had not been edited (LRRK2 +/+) or were knockout (LRRK2 -/-) were exposed to IFN-ɣ overnight or not treated (ctrl) then with or without 1mM LLOMe for fifteen minutes. After protein extraction, blots for LRRK2, pRAB10, pRAB12 and pRAB29 and appropriate controls were performed, in this case using b-actin (ACTB) as a protein reference. LRRK2 protein levels were increased by IFN-ɣ alone and, to a stronger extent by IFN-ɣ plus LLOMe (i), with similar effects on pRAB10 (j), pRAB12 (k) and pRAB29 (l) only in LRRK2 +/+ lines and not in LRRK2 -/- iMicroglia. *, *p*<0.05; ** *p* <0.001; ***, *p*<0.0001 by two-way ANOVA for treatment by genotype (For LRRK2 induction - F_3,16_ treatment = 76.84; F_1,16_ genotype = 255.54; F_3,16_ interaction = 76.84; For pRAB10 - F_3,16_ treatment = 133; F_1,16_ genotype = 252.2; F_3,16_ interaction = 132.4; For pRAB12 - F_3,16_ treatment = 37.06; F_1,16_ genotype = 112.8; F_3,16_ interaction = 37.03; For pRAB29 - F_3,16_ treatment = 13.83; F_1,16_ genotype = 86.98; F_3,16_ interaction = 12.43) with Tukey’s *post-hoc* test to compare all conditions. For clarity, only comparisons in the LRRK2+/+ cells are shown. For all blots, each lane is a replicate culture and molecular weight markers are shown on the right. For all graphs, bars show mean values, error bars are standard deviation, and individual points represent independent cultures. *N = 3 independent differentiations; representative of 3 independent experiments*.

To confirm that the observed effects on RAB phosphorylation are due to LRRK2 we repeated these experiments with isogenic LRRK2 knockout lines that we have previously described (27) which contain frameshift mutations at amino acid 1339 resulting in loss of functional LRRK2 protein. After overnight treatment with IFN-ɣ, we subsequently exposed cells for 15 minutes to 1mM L-Leucyl-L-Leucine methyl ester (LLOMe), which induces lysosomal membrane permeabilization and increases LRRK2 activity in various cell types (19) including in iMicroglia (17). We found that IFN-ɣ, but not LLOMe alone, significantly increased LRRK2 protein levels, with a combined effect of both treatments that was greater than IFN-ɣ alone (Fig. 1h, i). Of note, a ∼150kDa band was seen with IFN-ɣ that was diminished after combined IFN-ɣ plus LLOMe treatment and absent entirely in LRRK2 knockout cells, suggesting that lysosomal damage may prevent protein degradation of intact LRRK2. Importantly, phosphorylated RAB10 (Fig. 1j), RAB12 (Fig. 1k), and RAB29 (Fig. 1l) were all increased by IFN-ɣ and further enhanced by combined treatment with LLOMe, extending our observations to multiple RAB substrates. Microglia cells derived from the LRRK2 knockout line showed no phosphorylation of RAB proteins under any conditions (Fig. 1h-l), indicating that LRRK2 is required for inflammation-induced RAB phosphorylation.

To evaluate reproducibility of these effects, we measured LRRK2 induction after IFN-ɣ treatment in two additional iPSC lines derived from different donors, the fibroblast-derived male lines KOLF2.1J (28) and WTC11 (29) (Supplementary Fig. S1d-i). Across different experiments, LRRK2 protein was increased by ∼5-8 fold over baseline after IFN-ɣ treatment in iMicroglia from all tested donors. We also examined whether IFN-ɣ would elevate LRRK2 levels in 35-day-old forebrain neurons or dopaminergic neurons differentiated from the same iPSC lines (27), including the LRRK2 KO line as a negative control. We observed a ∼1.3-fold increase in LRRK2 levels in forebrain neurons that was absent from LRRK2 knockout lines (Supplementary Fig. S1j,k). LRRK2 expression was extremely low in dopaminergic neurons and was not altered by IFN-ɣ treatment (Supplementary Fig. S1j, k). These results demonstrate that LRRK2 protein expression and activity is robustly induced in iMicroglia to a much greater extent than neurons.

### Changes in chromatin accessibility in the LRRK2 promoter after interferon-ɣ treatment

Prior reports suggest that IFN-ɣ regulates LRRK2 at the level of mRNA in iPSC derived neurons (23). We therefore asked whether we would see the same in iMicroglia and whether the LRRK2 promoter on chr12 would respond to IFN-ɣ treatment by changes in chromatin structure. To do so, we used a multiome assays to simultaneously capture both the transcriptome and chromatin accessibility profiles of individual cells including LRRK2 wild type and knockout iMicroglia with or without IFN-ɣ (Fig. 2a). iMicroglia transcriptomes formed two distinct clusters defined by IFN-ɣ treatment on a UMAP projection, with LRRK2 mRNA expression being much higher in the IFN-ɣ treated subcluster (Fig. 2b). ATAC-seq data showing open chromatin showed the emergence of a new peak at ch12:40223500-40224000 (GRCh38 hg38 coordinates, labeled as ‘Peak A’ in Fig. 2c), an increase in the amplitude of two other peaks (Peak B, ch12:40220110-40220600, and Peak C, ch12:40209200-40209000, Fig. 2c).

**Figure 2.**
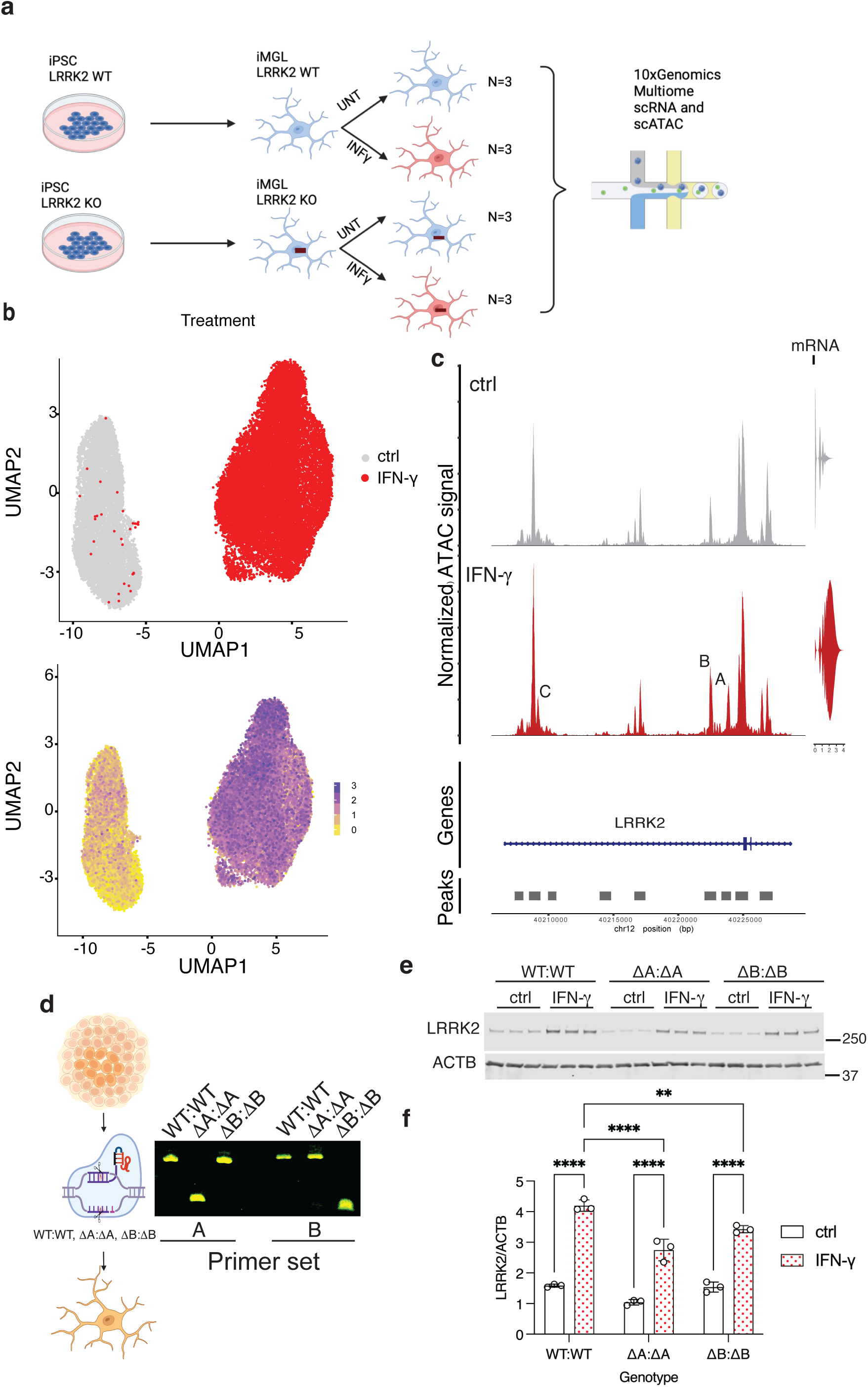
Changes in chromatin accessibility in the LRRK2 promoter after IFN-ɣ treatment. (a) Schematic showing LRRK2 WT and KO iMicroglia processed for Multiome scRNA-seq and scATAC-seq. (b) UMAP projections of iMicroglia scRNA-seq showing LRRK2 induction by IFN-ɣ. Each dot represents a single cell colored by treatment (ctrl or IFN-ɣ, upper panel) or by LRRK2 expression levels (lower panel, Scale on right). A total of 27, 406 cells are shown (c) Chromatin accessibility in the LRRK2 promoter region in iMicroglia without (ctrl, grey) or with IFN-ɣ (red) treatment, shown alongside a violin plot displaying the distribution of LRRK2 RNA expression in the same cells. Peak A (ch12:40223500-40224000) emerged after IFN-ɣ stimulation, and the intensity of Peaks B (ch12:40220110-40220600) and C (ch12:40209200-40209000) increased following IFN-ɣ stimulation. ‘TSS’ indicates the position of the LRRK2 transcription start site (TSS). (d) CRISPR/Cas9 editing of Peak A and Peak B in iPSCs, with PCR genotyping (right) showing positive homozygous (ΔA:ΔA) and homozygous (ΔB:ΔB) clones. (e) Western blot analysis of iMicroglia derived from WT, homozygous (ΔA:ΔA) Peak A deletion mutant, and homozygous (ΔB:ΔB) Peak B deletion mutant showing lower LRRK2 induction after IFN-ɣ treatment for both Peak A and B mutants, as well as decreased basal LRRK2 expression for Peak A mutant. Quantification of LRRK2 relative to ACTB are shown in (f), ** *p* <0.001; ***, *p*<0.0001 by two-way ANOVA for IFN-ɣ treatment by genotype (F_1,12_ treatment =544.7; F_2,12_ genotype = 42.64; F_2,12_ interaction = 9.704; n=3 replicate cultures per genotype and treatment) with Tukey’s *post-hoc* test to compare all conditions. For clarity, only comparisons for treatment effects in each line, and between lines for treated samples, are shown. Each lane is a replicate culture and molecular weight markers are shown on the right. All bars show mean values, error bars are standard deviation and individual points represent independent cultures. *N = 3 independent differentiations; representative of 3 independent experiments*.

These results imply that IFN-ɣ regulates LRRK2 via changes in the chromatin structure of the LRRK2 promoter. To test this idea, we generated CRISPR knockout clones lacking the peaks ‘A’ or ‘B’ (Fig. 2d and Supplementary Fig. S2). Deletion of peak A but not peak B decreased basal LRRK2 expression under untreated conditions and deletion of either of the peak regions resulted in diminished LRRK2 induction after IFN-ɣ treatment (Fig. 2e, f). Neither peak deletion resulted in complete loss of LRRK2 induction, suggesting that both contribute to regulation, possibly by co-operative mechanisms. These results show that the effects of IFN-ɣ on LRRK2 expression proceed via chromatin accessibility changes in the immediate promoter region, resulting in enhanced transcription of the LRRK2 gene.

### JAK/STAT signaling and ATF5 control LRRK2 expression after interferon-ɣ treatment

To identify signaling pathways that control LRRK2 expression after IFN-ɣ treatment, we analyzed pseudo-bulk single cell differential gene expression in the multiome dataset described in figure 2. We first confirmed that the effects of IFN-ɣ on ATAC-Seq data and LRRK2 mRNA expression were consistent in LRRK2 wild type and KO cells (Supplementary Fig. S3a-c). We next identified ∼5000 differentially expressed genes (DEGs) with greater than 2-fold change and an adjusted p < 0.05 in WT iMicroglia in response to IFN-ɣ treatment (Supplementary Fig. S3d). Of note, this strong transcriptional response to IFN-ɣ treatment was also seen in LRRK2 knockout iMicroglia (Supplementary Fig. S3e). There were ∼2000 genes each that were upregulated or downregulated concordantly in both genotypes (Supplementary Fig. S3f). Principal component analysis of the 500 most variable genes showed that treatment was associated with PC1 and ∼96% of variance, whereas genotype was associated with PC2, explaining only ∼3% of total variance (Supplementary Fig. S3g). Looking at most significant DEGs associated with IFN-ɣ treatment further confirmed that WT and KO cells responded similarly to the treatment (Supplementary Fig. S3h). Collectively, these results show that IFN-ɣ has, as expected, a profound effect on the transcriptional profile of iMicroglia but that LRRK2 has only a modest effect on how cells respond, demonstrating that LRRK2 is downstream of IFN-ɣ.

Next, to identify transcriptional control pathways involved in regulation of LRRK2, we selected 172 genes from the DEG list upregulated after IFN-ɣ treatment irrespective of LRRK2 genotype (Supplementary Fig. S3f) that are either transcription factors or involved in transcriptional regulation (Fig. 3a) based on a comprehensive human transcription factor gene catalog (30). We made an siRNA library to knock down each of these candidate transcription factors and TF regulators in iMicroglia under IFN-ɣ treatment and measured LRRK2 expression by qPCR. This secondary screen confirmed thirteen candidates that significantly affected LRRK2 expression after IFN-ɣ exposure (Fig. 3b). STRING analysis suggested that these molecules were linked to two interacting protein pathways: STAT1 and ATF5 (Fig. 3c). Based on this analysis, we further validated several linked candidate genes (ATF5, JAK1, JAK2, NDC80, NOD2, PIM1, STAT1, and TRIB3) using qPCR in independent experiments (Fig. 3d). Of note, NOD2 is a strong candidate gene for risk of Crohn’s disease at the Chr16 locus (31).

**Figure 3.**
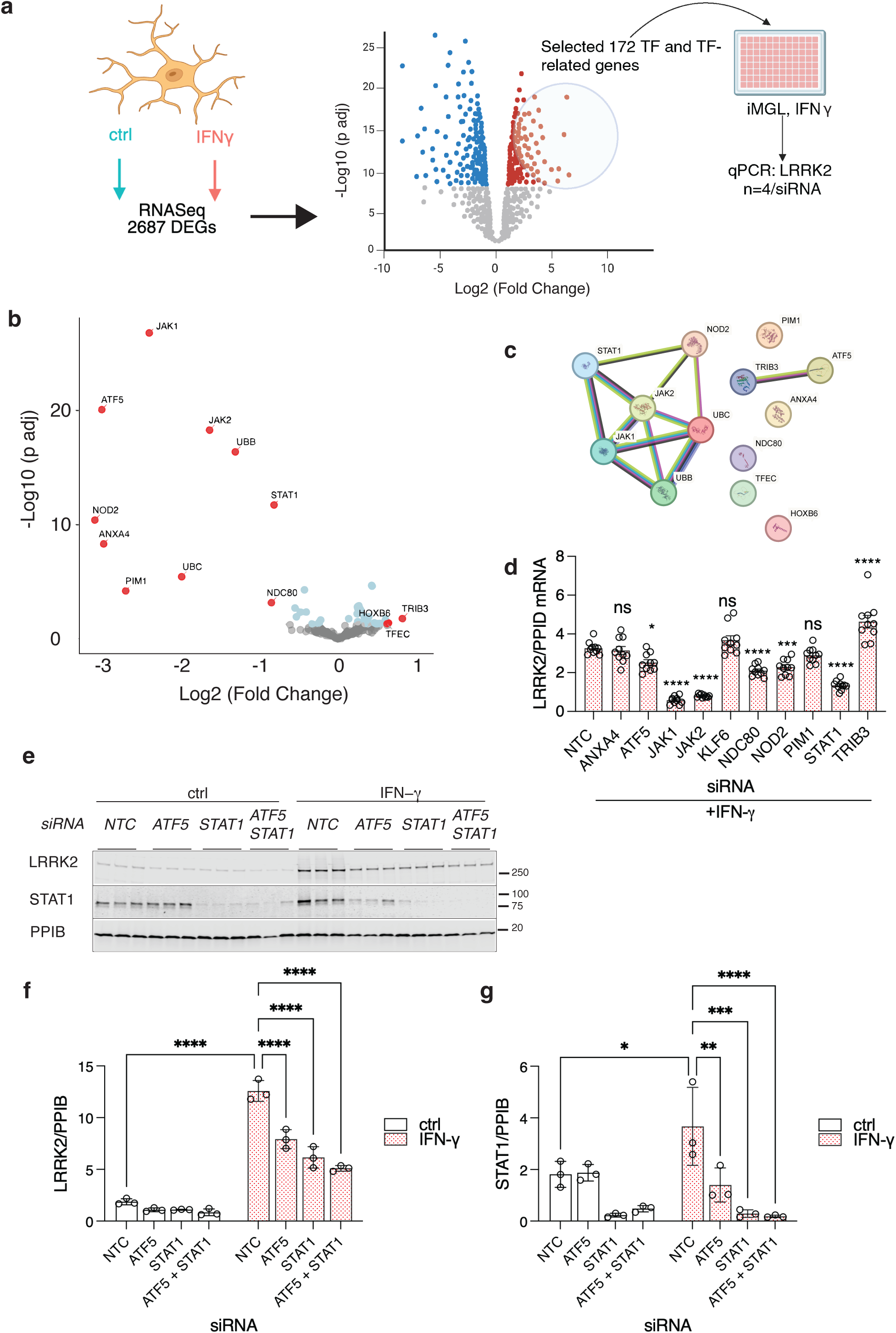
JAK/STAT signaling and ATF5 control LRRK2 expression after IFN-ɣ treatment. (a) Schematic of the screen. From 2,289 genes upregulated following IFN-ɣ treatment in iMicroglia (see Supplementary Fig S2), we selected 172 transcription factors (TFs) and related genes for a custom siRNA library to test whether knockdown of any of these genes decreased LRRK2 after IFN-ɣ treatment using qRT-PCR. (b) Volcano plot showing effects of each siRNA on LRRK2 comparing fold change compared to control siRNA. Thirteen genes that significantly affect LRRK2 mRNA levels with IFN-ɣ treatment are highlighted in red; FDR-adjusted p<0.05 using two-sided t-tests against control siRNA, with ≥1.5-fold change. (c) STRING database analysis of candidate genes revealed interactions between various members of the JAK/STAT signaling pathway, and a reported interaction between ATF5 and TRIB3. (d) Targets were further validated using qPCR in independent experiments to confirm decrease in LRRK2 mRNA. *, *p*<0.05; *** *p* <0.0001; ****, *p*<0.00001 by one-way ANOVA for siRNA (F_10,99_ = 59.69, n=10 replicate cultures per siRNA) with Dunnett’s *post-hoc* test to compare against non-targeting control (NTC) siRNA.(e) Western blot confirmed that there was a significant decrease in LRRK2 induction at the protein level when STAT1 and ATF5 were knocked down (f) as well as a decrease in STAT1 when ATF5 was knocked down. *, *p*<0.05; ** *p* <0.001; ***, *p*<0.0001 by two-way ANOVA for IFN-ɣ exposure by siRNA (LRRK2 - F_1,16_ treatment = 669; F_3,16_ siRNA = 51.87; F_3,16_ interaction = 30.66; STAT1 - F_1,16_ treatment = 1.273; F_3,16_ siRNA = 21.61; F_3,16_ interaction = 4.419) with Tukey’s *post-hoc* test to compare all conditions. For clarity, only comparisons for siRNA in the IFN-ɣ treated cells are shown as well as baseline induction. On the blot, each lane is a replicate culture and molecular weight markers are shown on the right. For all graphs, bars show mean values, error bars are standard deviation and individual points represent independent cultures. *N = 3 independent differentiations; representative of 3 independent experiments*.

Additionally, we confirmed that there was a significant decrease in LRRK2 induction at the protein level when STAT1 and ATF5 were knocked down (Fig. 3e, f). Knockdown of both transcription factors were effective in blocking IFN-ɣ induced LRRK2 protein levels. We could not identify an antibody that recognizes ATF5, but we did see a notable decrease in STAT1 levels with ATF5 knockdown (Fig. 3g). This suggests that ATF5 regulates the expression of STAT1 in these cells, consistent with analysis of the STAT1 promoter and possibly related to the known role of ATF5 in regulating stress responses (32).

We also identified two kinases, JAK1 and JAK2, which are known to phosphorylate and activate STAT1. We knocked down these kinases both individually and together, showing a significant reduction in LRRK2 induction, particularly when both kinases were knocked down (Supplementary Fig. S4a-e). Additionally, we confirmed that the Tyr701 phosphorylation site on STAT1 (24) is crucial for STAT1 activation and LRRK2 induction (Supplementary Fig. S4a, d). Interestingly, the knockdown of JAK1 and JAK2 also affected overall STAT1 levels as well as phosphorylation (Supplementary Fig. S4a).

IRF1 is a STAT1-target gene involved in secondary group-I and group-II interferon responses by activating the transcription of genes stimulated by IFN-ɣ. To determine whether STAT1 activation is directly responsible for LRRK2 induction or if it is mediated through secondary IRF1 activation, we examined the effects of IRF1 knockdown. We found that knocking down IRF1 did not influence LRRK2 induction (Supplementary Fig. S4 f-l), suggesting that STAT1 activation directly drives the increased expression of LRRK2 in response to IFN-ɣ rather than indirectly through IRF1. Overall, the results from our siRNA screen and validation experiments suggest that LRRK2 regulation by IFN-ɣ proceeds through the JAK-STAT pathway, with additional regulation by ATF5 but not IRF1.

### STAT1 binding to the LRRK2 promoter is associated with regulation of long non-coding RNAs that stabilize LRRK2 expression

Based on the above observations, we considered if STAT1 binds the LRRK2 promoter directly. We performed ChIP-Seq with STAT1 antibodies under control and IFN-ɣ conditions and compared these with our previously generated ATAC-seq data (Fig. 4a). The strongest STAT1 binding site overlaps with ATAC-seq peak ‘B’ that was shown to be enhanced after IFN-ɣ exposure. These results support the hypothesis that STAT1 directly binds the LRRK2 promoter.

**Figure 4.**
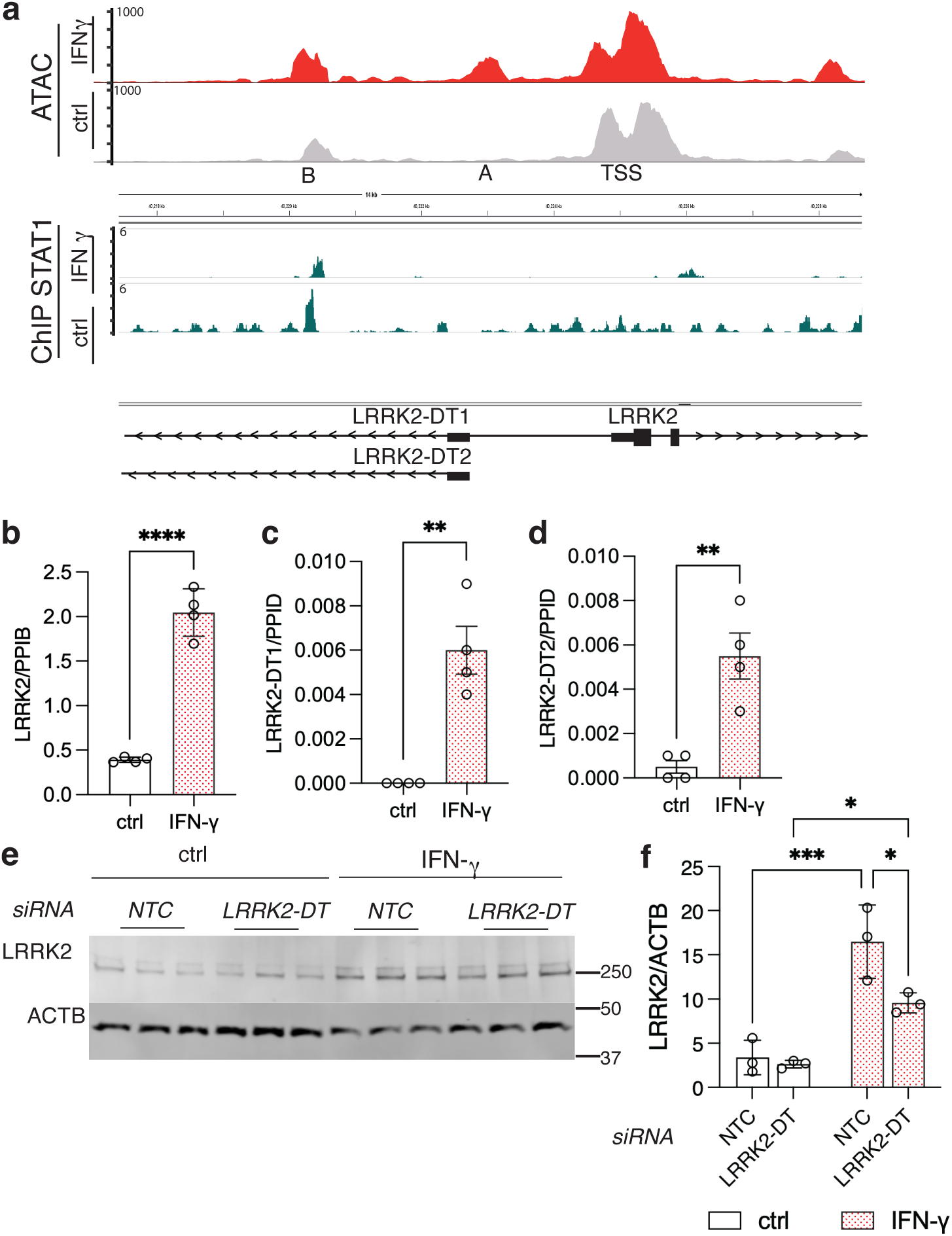
STAT1 binds to the LRRK2 promoter, and LRRK2 expression is associated with the regulation of long non-coding RNAs. (a) LRRK2 promoter region showing ATAC-seq (upper panels) and ChIP-seq peaks (lower panels) under control (grey) and IFN-ɣ (red) treatment. The strongest STAT1 binding site overlaps with ATAC-seq peak ‘B,’ which was shown to be enhanced after IFN-ɣ exposure. ATAC-seq peak ‘A’ overlaps the promoter region shared by LRRK2 and two annotated long non-coding RNAs, LRRK2-DT1 and LRRK2-DT2. (b-d). qRT-PCR analysis of iMicroglia treated with or without IFN-ɣ showed a significant increase not only in LRRK2 mRNA (b; ****, *p*<0.001 by unpaired t-test, t=12.32, df=6, n=4 replicate cultures per treatment), but also in LRRK2-DT1 (c; **, *p*=0.0014 by unpaired t-test, t=5.56, df=6, n=4 replicate cultures per treatment) and LRRK2-DT2.(d; **, *p*=0.0036 by unpairest t-test, t=4.629, df=6, n=4 replicate cultures per treatment). (e) Western blot analysis demonstrated a decrease in LRRK2 induction in iMicroglia treated with LRRK2-DT siRNAs compared to the non-targeting control (NTC). Quantification of LRRK2 relative to ACTB is shown in (f). *, *p*<0.05; ** *p* <0.001; ***, *p*<0.0001 by two-way ANOVA for IFN-ɣ exposure by siRNA (F_1,8_ treatment = 53.3; F_1,8_ siRNA = 7.86; F_1,8_ interaction = 5.06; n=3 replicate cultures per) with Tukey’s *post-hoc* test to compare all conditions. For all blots, each lane is a replicate culture and molecular weight markers are shown on the right. For all graphs, bars show mean values, error bars are standard deviation and individual points represent independent cultures. *N = 3 independent differentiations; representative of 3 independent experiments*.

During this analysis, we also observed that the ATAC-Seq peak A and STAT1-binding region (peak B) are adjacent to two annotated long non-coding RNAs, LRRK2-DT1 and LRRK2-DT2. Using qRT-PCR, we were able to show that both long non-coding transcripts were elevated 24 hours after IFN-γ treatment (Fig. 4b-d). We measured the time course of increases in LRRK2 and LRRK2-DT mRNA after IFN-ɣ treatment.

LRRK2-DT levels showed a significant increase only after 24 hours (Supplementary Fig. S5a, b) whereas LRRK2 expression was significantly elevated as early as 2 hours post-treatment (Supplementary Fig. S5c).

Long non-coding RNAs (lncRNAs) are generally expressed at much lower levels than mRNAs that they regulate. Recent literature (33) suggests that the low expression of lncRNAs can be critical to ensure that they are directed to specifically regulate their target mRNAs. We noted that LRRK2-DT mRNA levels were approximately 100 times lower than LRRK2 mRNA levels, suggesting that LRRK2-DT might affect LRRK2 induction. Consistent with this hypothesis, LRRK2-DT knockdown limited LRRK2 induction at the protein level following IFN-ɣ treatment (Fig. 4e, f). These observations show that there is coregulation of long non–coding RNA and mRNA for LRRK2, leading to an overall stabilization of LRRK2 levels, likely secondary to initial regulation by STAT1.

### LRRK2 is not detectably induced by interferon-ɣ in mouse brain

To evaluate whether we could recapitulate LRRK2 induction *in vivo*, we administered mouse Ifn-ɣ into the substantia nigra (SN) and striatum (ST) of the Lrrk2 wild type and knockout mice, with PBS as a vehicle control (Fig. 5a). Of note, the Lrrk2 knockout has a deletion of exon 2 that leads to a premature stop codon in exon 3 and thus has Lrrk2 mRNA but no functional protein. After dissection of individual regions, we performed single cell multiome to examine LRRK2 expression across cell types. Based on known cell markers (Fig. 5b), UMAP projections of the single cell data identified all expected types of cells in the substantia nigra (Fig. 5c). Ifn-ɣ had profound effects on the transcriptome of several cell types, notably glial cells that could be readily distinguished by treatment on UMAP (Fig. 5d). A similar separation by known cell markers was obtained in the striatum where a clear response to Ifn-ɣ was also seen (Supplementary Fig. S6a-c). We found many differentially expressed genes between PBS and Ifn-ɣ treated samples. It is known that Ifn-ɣ exposure induces Stat1 expression, which we confirmed in our data for all cell types in the substantia nigra (Fig. 5e) and the striatum (Supplementary Fig. S6d) in both Lrrk2 wild type and knockout mice. Despite this positive confirmation that all animals mounted a strong response to Ifn-ɣ, we did not see any change in expression of Lrrk2 after Ifn-ɣ exposure in the mouse *in vivo* in either substantia nigra (Fig. 5f) or striatum (Supplementary Fig. S6e). These show that Ifn-ɣ does not cause Lrrk2 induction in microglia in the intact mouse brain, despite proven engagement of immune signaling.

**Figure 5.**
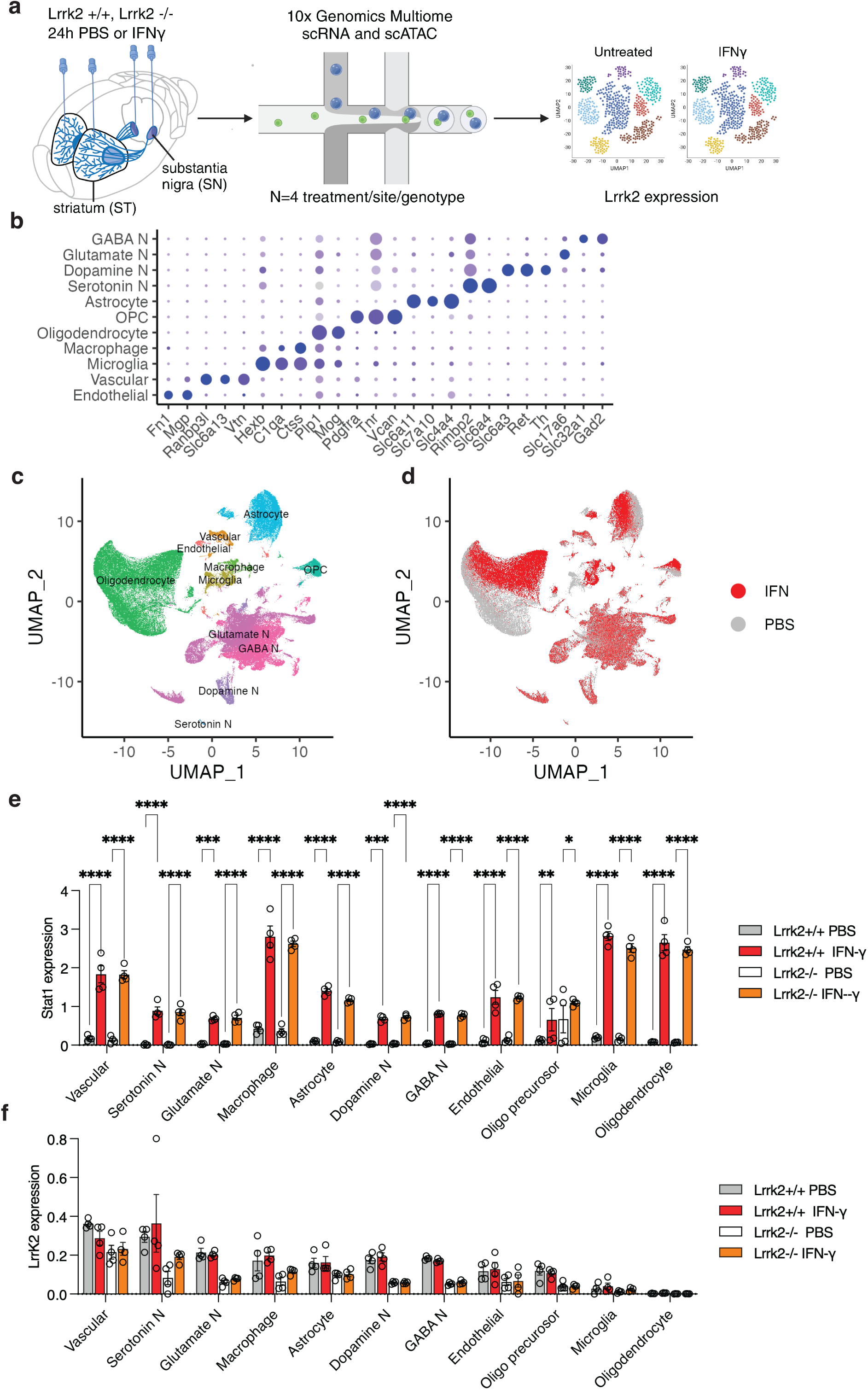
Interferon-γ does not significantly induce LRRK2 expression in the mouse substantia nigra. (a) Experimental design for assessing IFN-ɣ induced *Lrrk2* expression in Lrrk2 wild type and knockout mouse brain regions. Mice were bilaterally injected with either PBS (as a control) or recombinant mouse IFN-γ into the striatum (ST) or substantia nigra (SN). After 24 hours, the injected regions were dissected and processed using multiome RNA-seq and ATAC-seq. (b) Dot plot analysis showing major cell subtypes in substantia nigra. The average expression level of a gene within a cell type, indicated by color intensity, and the proportion of cells within that type expressing the gene, represented by dot size. (c,d) UMAP visualization of major cell populations (148,542 cells) in the mouse substantia nigra colored by identified cell types (c) or treatment (d). (e) STAT1 expression across major cell types in the substantia nigra, comparing ± Ifn-ɣ treatment and LRRK2 wild-type (WT) vs. knockout (KO) mice. (f) LRRK2 expression across major cell types in the substantia nigra, comparing ± Ifn-ɣ treatment and LRRK2 wild-type (WT) vs. knockout (KO) mice. \**p*<0.05; ** *p* <0.001; ***, *p*<0.0001 by three-way ANOVA for Ifn-ɣ exposure by genotype and by cell-type (For Stat1 - F_10,132_ for cell type = 68.95; F_1,132_ for genotype = 0.02187; F_1,132_ for IFN-ɣ exposure = 1718; For Lrrk2 - F_10,132_ for cell type = 28.29; F_1,132_ for genotype = 99.33; F_1,132_ for IFN-ɣ exposure = 1.713; n=4 mice per genotype and treatment group) with Tukey’s *post-hoc* test for individual comparisons; for clarity, only comparisons between PBS and IFN-ɣ for each cell type that were significant (p<0.05) are shown.

### LRRK2 induction by interferon-ɣ is muted in murine compared to human cells of the monocyte lineage

The lack of induction of Lrrk2 by Ifn-ɣ in the mouse brain may be related to multiple factors including *in vitro* compared to *in vivo* systems, sex differences, species differences or other unknown factors. To address these different possibilities, we first examined murine glial cell cultures, which we have previously shown contain active Lrrk2 (19). Despite robust induction of Stat1 by mouse Ifn-ɣ, we were unable to detect an increase in mouse Lrrk2 (Supplementary Fig. S7a-c). Given the robust induction of LRRK2 in human iMicroglia, this data suggests that there is a species difference between human and mouse LRRK2 orthologs in their sensitivity to interferons. However, iMicroglia may not be perfectly analogous to brain microglia in terms of transcriptional control.

Furthermore, the potential role of LRRK2 in both CNS and peripheral diseases indicate that both immune cells within or outside the brain might express LRRK2 and be subject to regulation by inflammatory signaling. We therefore looked for a system where we could as closely match cell types across species to examine how interferon regulates LRRK2. We chose bone marrow derived cells that we could acquire commercially from human donors or obtain from mice in the laboratory and could differentiate into monocytes, macrophages or dendritic cells based on growth factor exposure (Fig 6a). We found that IFN-ɣ treatment of human monocytes (Fig. 6b, e,f), macrophages (Fig. 6 c, g, h) or dendritic cells (Fig. 6d, i, j) all induced robust increases in LRRK2 expression (Fig. 6e, g, i) and pRAB10 (Fig. 6f, h, j). In contrast, while we were able to demonstrate induction of Lrrk2 in murine monocytes (Fig. 6k, n), there was no significant increase in pRab10 (Fig. 6o). Additionally, we were unable to find any increases in Lrrk2 or pRab10 in mouse macrophages (Fig 6l, p,q) or dendritic cells (Fig. 6m, r, s). Even when we saw evidence of induction of Lrrk2 ex pression in mouse monocytes, the effects were more modest than seen in the equivalent human cells. For example, at 48h of treatment of human monocytes, LRRK2 was induced by ∼12 fold (range 10 to 15-fold) whereas in mouse monocytes the induction was ∼8 fold (range 7.3 to 9.3-fold). Similarly, pRAB10/RAB10 in human monocytes was ∼4.3 fold (range 3.7 to 4.6-fold) but in mouse cells was ∼2 fold (range 1.8 to 2.1-fold). In all cases, there was a clear accumulation of STAT1/Stat1, confirming all cell types responded to interferon signaling. Consistent effects were seen (or not) in samples from additional human donors or individual mice (Supplementary Fig. S7d-u). These results suggest that there may be fundamental differences between how human LRRK2 and mouse Lrrk2 are regulated by interferon signaling that is likely additive to the previously reported lower kinase activity of the murine orthologue (34).

**Figure 6.**
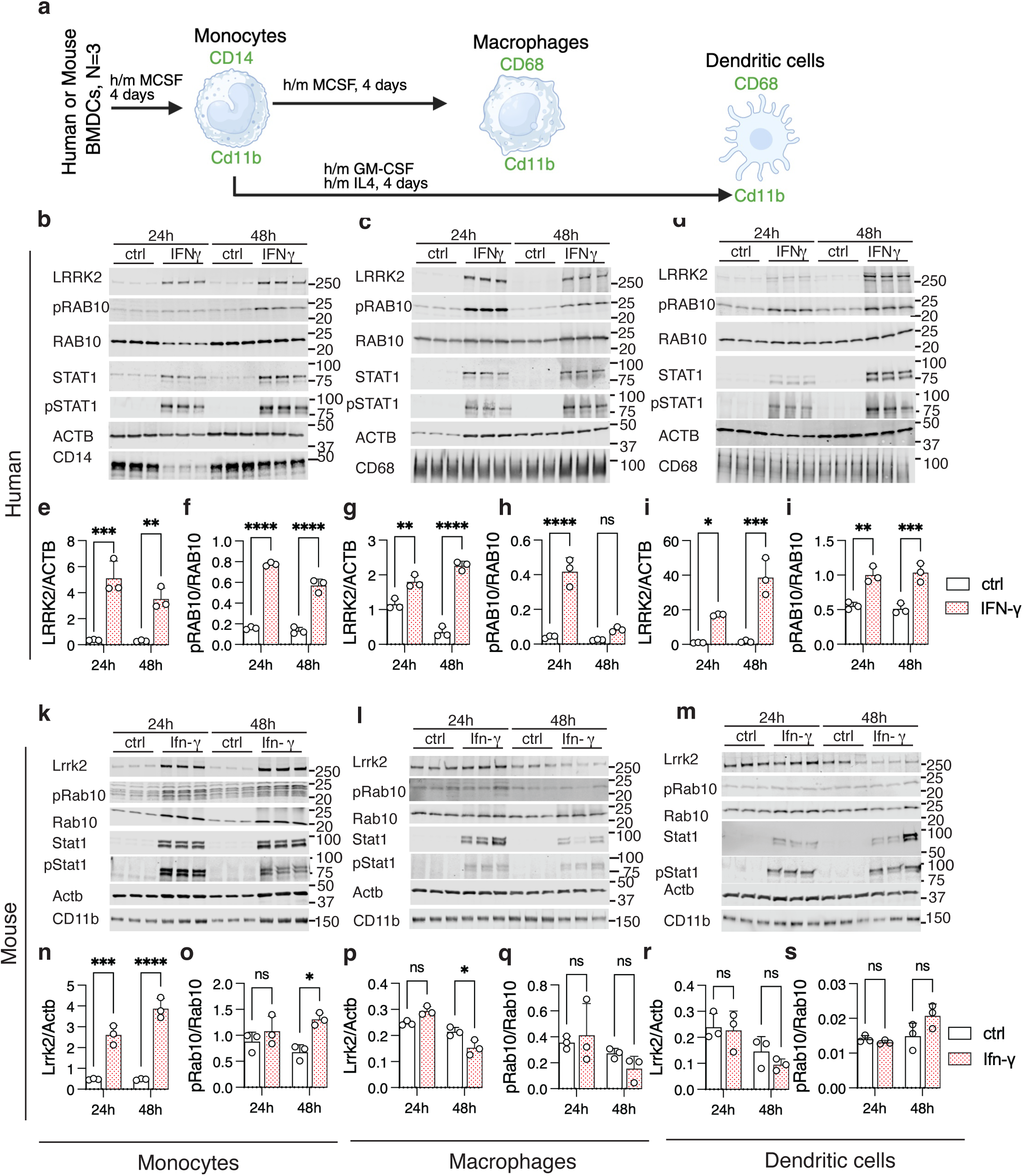
Interferon-γ induces more modest LRRK2 expression in murine compared to human monocyte-lineage cells. (a) Diagram showing the differentiation of monocytes, macrophages, and dendritic cells from human and mouse bone marrow–derived cells (BMDCs). Panels (b–d) show representative western blots for human monocytes (b), macrophages (c), and dendritic cells (DCs) (d) respectively, for LRRK2, pRAB10, RAB10, STAT1, pSTAT1 (Y701), ACTB, CD14, CD68, and CD11b protein levels after 24 and 48 hours of IFN-ɣ treatment. Quantification of protein expression is shown in human monocytes (e, LRRK2; F_1,8_ treatment = 78.4; f, pRAB10; F_1,8_ treatment = 607.9), macrophages (g, LRRK2; F_1,8_ treatment = 208.6; h, pRAB10; F_1,8_ treatment = 85.9) and dendritic cells (i, LRRK2; F_1,8_ treatment = 72.6; j, RAB10; F_1,8_ treatment = 70.5). Panels (k–m) show representative western blots for mouse monocytes (k), macrophages (l), and DCs (m), respectively. Quantification of protein expression is shown in mouse monocytes (n, Lrrk2; F_1,8_ treatment = 78.4; o, pRab10; F_1,8_ treatment = 13.96), macrophages (p, Lrrk2; F_1,8_ treatment = 0.414; q, pRab10; F_1,8_ treatment = 0.14) and dendritic cells (r, Lrrk2; F_1,8_ treatment = 1.01; s, pRab10; F_1,8_ treatment = 2.73). In all graphs, ns p>0.05, \**p* < 0.05; **p < 0.001; \*\*\**p* < 0.0001 by two-way ANOVA assessing treatment and time point (values of F given above), followed by Tukey’s post-hoc test for pairwise comparisons. For clarity, only comparisons between control and interferon exposure are shown within each time point using n=3 replicate cultures per treatment and time point.

### LRRK2 is induced by interferon-γ in human brain slices *ex vivo*

As a model to test whether we could induce LRRK2 expression in the human brain, we obtained freshly dissected brains from two surgical resection cases and acutely cultured slices for ∼6h *ex vivo* without or with exposure to IFN-ɣ (Fig. 7a). Projecting single cell RNA-seq data on UMAP, we were able to show that both donors (Fig. 7b) and treatment types (Fig. 7c) contained major cell types of the brain (Fig. 7d) based on known cell markers (Supplementary Fig. S8a). Oligodendrocytes represented a majority of recovered cells in these samples (Supplementary Fig. S8b) and quality control metrics for single cell RNA-Seq were consistent across cell types and treatments (Supplementary Fig. S8c-e).

**Figure 7.**
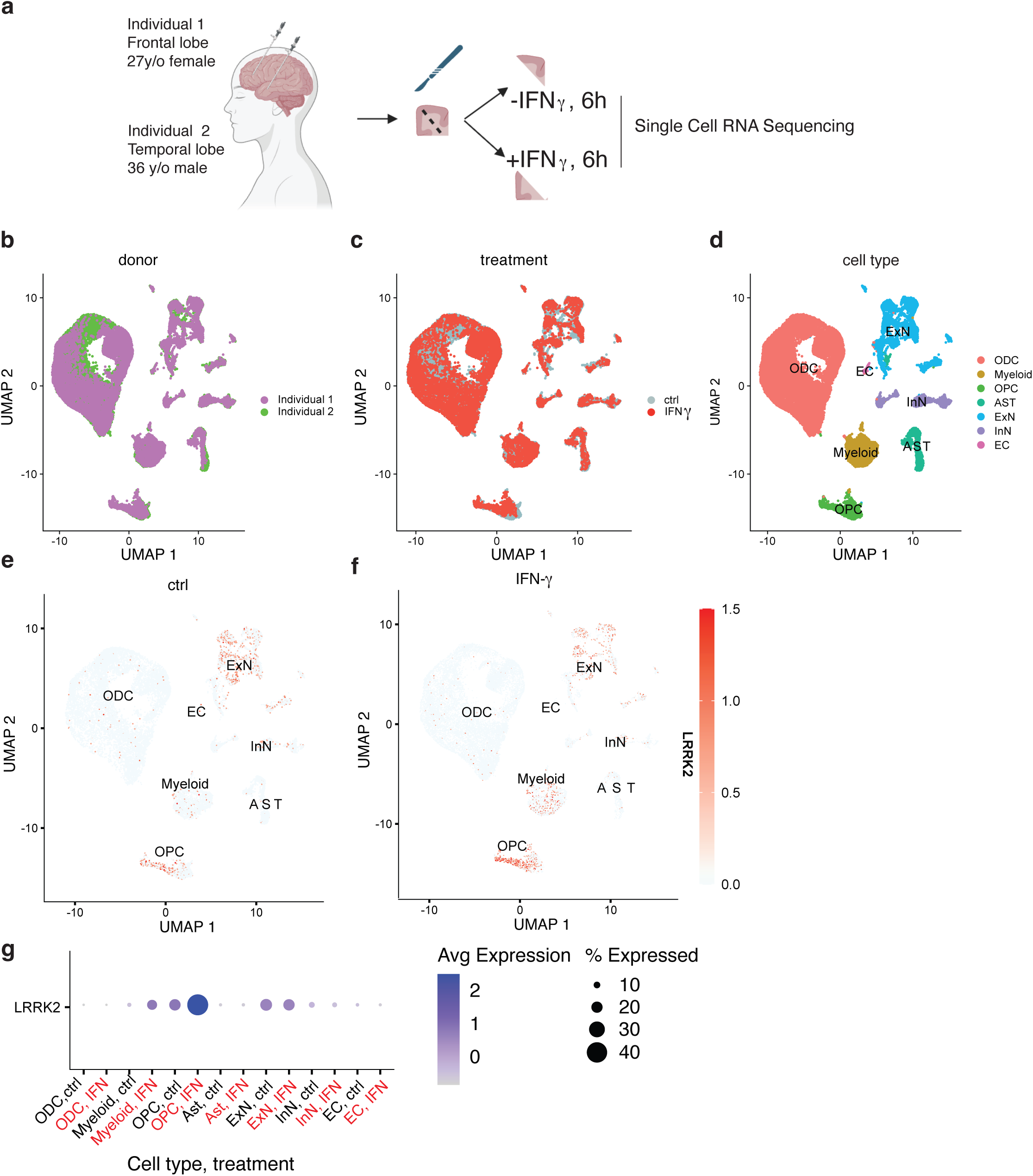
LRRK2 is induced by interferon-γ in human brain slices *ex vivo*. (a) Diagram showing the experimental outline using freshly dissected brain tissue from two surgical resection cases. Brain slices derived from frontal lobe (Individual 1) and temporal lobe (Individual 2) were acutely cultured *ex vivo* for six hours, without or with exposure to IFN-ɣ and processed for single cell RNA sequencing. (b) Clustering UMAP plots from the human brain sections showing cell distributions based on (a) individual, (c) IFN-ɣ treatment, and (d) major cell types. (e,f) Feature plot showing *LRRK2* expression without or with IFN-ɣ treatment. The color bar indicates relative expression levels. (g) *LRRK2* expression across major cell types in human brain sections, comparing conditions with and without IFN-ɣ treatment, colored by expression levels and sized by percentage of cells that express LRRK2 within each cell type. ODC, oligodendrocytes; OPC, oligodendrocyte precursor cells; Ast, astrocytes; ExN, excitatory neurons; InN, inhibitory neurons; EC, endothelial cells. A total of 36,526 cells were used for analysis: 18,440 cells from control samples and 18,086 cells from IFN-γ–treated samples; Individual 1 (17,946 cells), Individual 2 (18,580 cells).

Under baseline conditions, LRRK2 was expressed in subsets of excitatory neurons and oligodendrocyte precursor cells (OPC, Fig. 7e), similar to prior data from the human brain (*17*). However, there was a clear increase in the percentage of microglia that expressed LRRK2 after IFN-ɣ exposure (Fig. 7f) Additionally, while expression of LRRK2 in microglia and oligodendrocyte precursor cells was higher after IFN-ɣ treatment, LRRK2 did not show a clear increase in other cells, including excitatory neurons (Fig. 7g). We also confirmed that we saw similar patterns of LRRK2 induction in samples from both donors that were available to us (Supplementary Fig. S8f,g). These results demonstrate that human microglia can respond to IFN-ɣ with upregulation of LRRK2 in the context of *ex vivo* exposure.

### LRRK2 inducibility by interferon-γ is enhanced by human-specific promoter elements

Based on the above results, we considered whether regulation of LRRK2 by IFN-ɣ may be more facile in human vs mouse cells due to the structure of the human LRRK2 promoter. We first performed a comparative genomic analysis of the regions upstream of LRRK2 that showed altered chromatin accessibility following IFN-ɣ treatment (peaks A-C in Fig. 2c). We found that two of these genomic elements, peak ‘A’ and peak ‘C’, showed moderate homology across species (Fig. 8a). In contrast, peak ‘B’, which we also nominated as being altered in organization and also bound STAT1 directly, was mostly present across the infraorder Anthropoidea (also referred to as Simiiformes, including humans, nonhuman apes, and African/American monkeys) but absent in all other species examined, including mice and other rodents (Fig. 8a). Of particular note, this region includes three predicted STAT1 binding sites (Fig. 8a). These results suggest a mechanism by which upregulation of LRRK2 by IFN-ɣ might be stronger in humans and other anthropoid primates compared to rodents, namely, genomically encoded elements that are responsive to JAK/STAT signaling.

**Figure 8.**
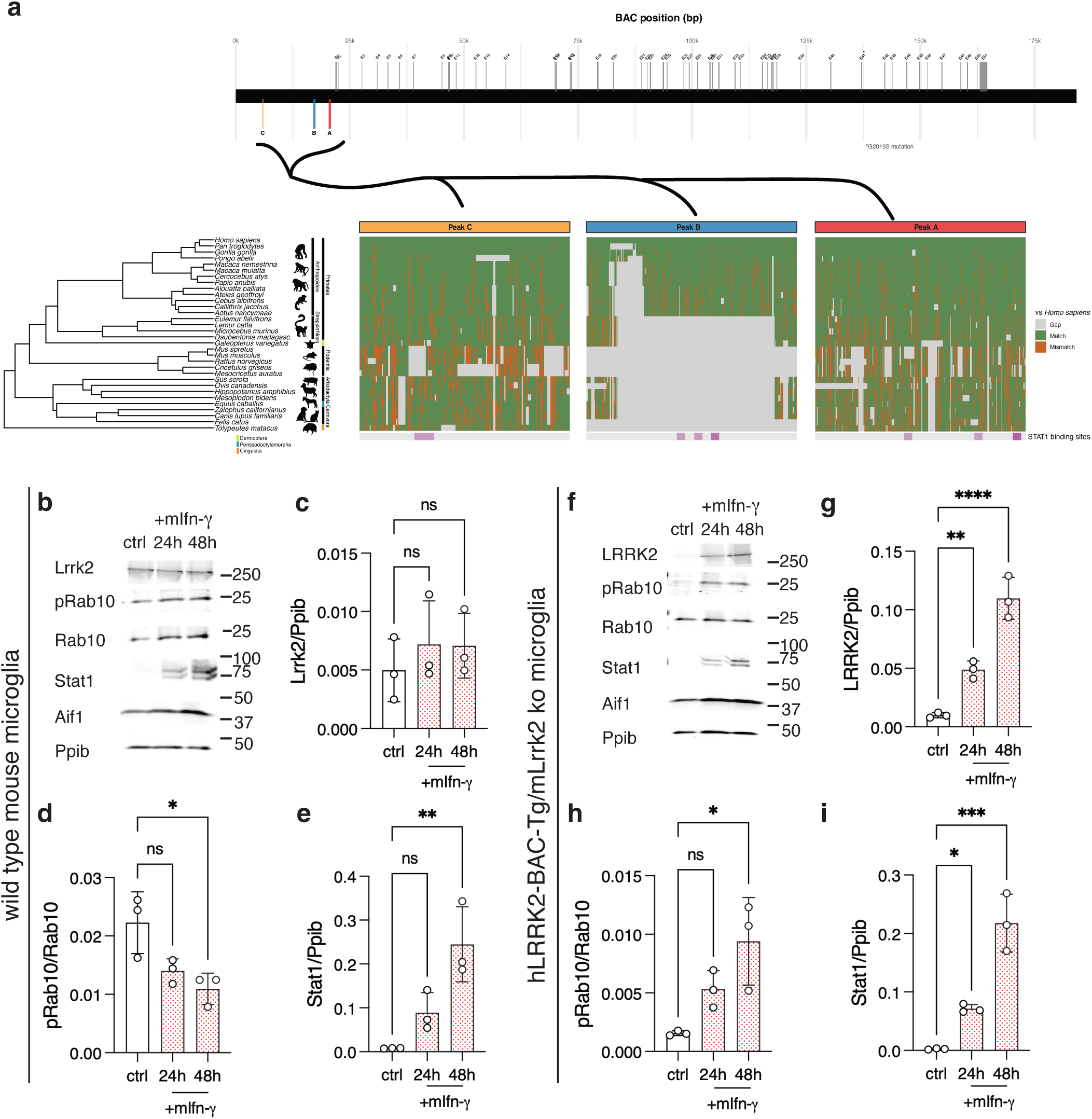
Human LRRK2 promoter elements are sufficient to confer interferon-γ inducibility to mouse microglia. (a) Comparative analysis of the LRRK2 promoter region across mammals.Top: human LRRK2 coding and promoter region with locations of exons (grey) and promoter peaks A (red), B (blue), and C (orange). Colors of peaks correspond to those in the bottom panel. Bottom (from left to right): Phylogenetic tree of select species (from TimeTree, see Methods). Species names are in italics. Humans (*Homo sapiens*; hg38) are at the top. The Orders represented in the tree (e.g., Primates, Rodentia) are named to the right of the tree and are indicated by black or colored blocks. Within Primates, the Suborder Strepsirrhini and Infraorder Anthropoidea are also shown. Icons of representatives from each clade are shown for reference (images from phylopic.org). Alignments to the human sequence for each peak sequence are shown: grey = gap, green = match, orange = mismatch. STAT1 binding sites are shown below each peak alignment plot in purple. (b-e) Primary mouse microglia from wild type animals were left untreated (ctrl) or treated with mouse interferon-gamma (mIfn-ɣ) for 24 or 48 hours as indicated then blotted for Lrrk2, pRab10, total Rab10, Stat1 as well as the microglial marker Aif1 and reference protein cyclophilin b (Ppib). Quantification of protein expression is shown for Lrrk2 (c; F_2,6_ = 0.48), pRab10 (d; F_2,6_ = 0.414) and Stat1 (e; F_2,6_ = 13.9). (f-i) Primary mouse microglia from Lrrk2 knockout animals that were also transgenic for human LRRK2 BAC (shown in a) were left untreated (ctrl) or treated with mouse interferon-gamma (mIfn-ɣ) for 24 or 48 hours as indicated then blotted for LRRK2, pRab10, total Rab10, Stat1 as well as the microglial marker Aif1 and reference protein cyclophilin b (Ppib). Quantification of protein expression is shown for LRRK2 (g; F_2,6_ = 60.1), pRab10 (h; F_2,6_ = 8.4) and Stat1 (i; F_2,6_ = 43.5). In all graphs; ns p>0.05; \**p* < 0.05; **p < 0.001; \*\*\**p* < 0.0001 by two-way ANOVA assessing treatment and time point (values of F given above), followed by Tukey’s post-hoc test for pairwise comparisons. For clarity, only comparisons between control and the two timepoints are shown using n=3 replicate cultures. For all blots, each lane is a replicate culture and molecular weight markers are shown on the right. For all graphs, bars show mean values, error bars are standard deviation and individual points represent independent cultures.

We next tested whether the human LRRK2 promoter would be sufficient to confer expression and IFN-ɣ sensitivity in microglia. We obtained a previously developed transgenic mouse where a ∼188kB BAC containing the human LRRK2 coding sequence plus upstream promoter region was integrated into mouse chr2 (C57BL/6J-Tg(LRRK2*G2019S)2AMjff/J, JAX stock #018785, referred to here as hLRRK2-BAC-Tg, Fig. 8a). We then crossed this BAC transgenic line to the Lrrk2 knockout mice line that was used in figure 5, thus giving only human LRRK2 expression from its endogenous genomic control elements but in a murine background. We separately purified microglia from wild type or hLRRK2-BAC-Tg/Lrrk2 ko mice, treated with murine IFN-ɣ and measured LRRK2 induction. Microglia from wild type mice responded to mouse IFN-ɣ did not show significant induction of Lrrk2 or pRab10 accumulation, despite robust induction of Stat1 (Fig. 8b-e). In contrast, microglia from the hLRRK2-BAC-Tg/Lrrk2 ko animals showed induction of LRRK2 and pRab10 over 24-48h, with associated Stat1 expression (Fig. 8f-i). Additionally, because the hLRRK2-BAG-Tg is present in the heterozygous state, and to allow direct comparison of LRRK2 levels, we repeated these experiments including microglia from Lrrk2 heterozygous (Lrrk2 +/-), hLRRK2-BAC-Tg/Lrrk2 ko, and wild-type (Lrrk2 +/+) mice (Supplementary Fig. S9a). We found that LRRK2 expression of the human BAC transgene was similar to that seen from mouse Lrrk2 in these cells but that there was profound induction of LRRK2 only in the cells form mice that were hLRRK2-BAG-Tg, despite similar Stat1 induction (Supplementary. Fog S9b,c). These results demonstrate that the proximal promoter region of human LRRK2, specifically regions that bind STAT1 and show rearrangement of chromatin accessibility after IFN-ɣ exposure, are required for induction of LRRK2 expression by IFN-ɣ, thus providing a mechanistic explanation for the observed species differences in LRRK2 regulation in human vs mice.

We note that the BAC transgene used in the above experiments contains that LRRK2 p.G2019S mutation associated with human PD. We considered whether this amino acid-changing variant would influence induction of LRRK2 by IFN-ɣ. We therefore took isogenic iPSC lines that were engineered to be either wild type, heterozygous (WT; GS) or homozygous (GS; GS) for the endogenous p.G2019S allele, differentiated to iMicroglia and treated with IFN-ɣ as above then blotted for relevant protein markers (Supplementary Fig. S9d). LRRK2 was reliably induced in all three iPSC lines (Supplementary Fig. S9e), to between 5 and 7 fold relative to baseline levels (range 3.9-8.3 fold). Notably, pRAB10 induction was similar between lines (Supplementary Fig. S9f), which is consistent with prior data that G2019S has only modest impact on pRAB10 (41). However, we did see a stronger effect of the G2019S mutation on pRAB12 and pRAB29 levels (Supplementary Fig. S9g,h). While the mechanistic basis for how different LRRK2 mutations phosphorylate RAB proteins to varying extents (41) is not clear, these results collectively show that the the LRRK2 p.G2019S allele does not impact strength of LRRK2 protein induction by IFN-ɣ.

## Discussion

Human aging is a complex set of events that are associated with multiple different hallmarks that include chronic inflammation (3,4). Illustrating the complex nature of the association between inflammation and age-related diseases, genes identified in PD also overlap with pathways in inflammatory disorders including CD (10,11,24). This suggests shared biological mechanisms between CNS and peripheral diseases where inflammation is an important precipitating event. In this context, determining how genetic risk factors that play causal but non-deterministic roles in age-associated diseases are regulated by inflammatory signaling is critical for understanding, modeling and intervening in disease processes.

In this context, we here describe in detail how LRRK2, associated with PD and CD as ostensibly distinct CNS and peripheral diseases, is regulated by IFN-ɣ. We show that LRRK2 is regulated by several related mechanisms in human iMicroglia that include JAK/STAT signaling, augmented by the stress-induced transcription factor ATF5 and two long non-coding RNA transcripts at the LRRK2 locus, leading to rearrangement of chromatin in the immediate promoter for LRRK2 and upregulation of LRRK2 mRNA levels. Surprisingly, this mechanism is not fully conserved in mice, where peripheral immune cells show increased expression of LRRK2 but central microglia do not. We show that these differences between species are genomically encoded due to STAT1 binding sites that arise in mammals between lemurs and higher infraorder primates. Importantly, reintroduction of human LRRK2 promoter elements into the mouse genome is sufficient to infer responsiveness to IFN-ɣ signaling. Of note, examination of a previously published (*35*) expression dataset also showed that Lrrk2 levels were higher in primates compared to all other mammalian species, while Jak1/2 and the homologous Lrrk1 protein did not shown this distinction (Supplementary Fig. S10a). Collectively, these data show that there are species-dependent mechanisms by which LRRK2 is regulated that are notably relevant to CNS-derived immune cells (Summarized in Supplementary Fig. S10b).

It is likely that upregulation of LRRK2 by IFN-ɣ signaling is functionally important in human microglia. Many studies have shown that, by phosphorylating RAB proteins, LRRK2 plays important roles in maintenance of lysosomal integrity under conditions of lysosomal membrane damage (36). There are many events related to inflammatory signaling that result in altered lysosomal function. For example, termination of cGAS-STING signaling requires lysosomal protein degradation (37,38). Equally, temporary lysosomal membrane permeabilization is important for cross-presentation of antigens between immune cells (39). It is notable that such transient events are appropriate if rapidly contained but maybe be maladaptive if not resolved quickly. Based on data presented here, we do not find evidence that LRRK2 modifies IFN-ɣ signaling per se, based on unaltered transcriptional outcomes between LRRK2 knockout and unmodified cells and animals. Rather, we postulate that LRRK2 is important for minimizing lysosomal damage consequent to inflammatory signaling.

The functional requirement for a species difference between human and mouse LRRK2 is less clear. Our comparative analysis suggests that the acquisition of crucial promoter elements that confer STAT1 responsiveness occurred during evolution of the primate lineage, likely between ancestral groups separating strepsirrhini and haplorhines. This is consistent with prior data (35) that identified microglial LRRK2 expression in humans and non-human primates but not artiodactyla or rodents. Of note, this species distinction is not seen with the homologous LRRK1 gene that appears to be basally active and phosphorylates RAB7. Geirsdottir et al (35) suggest that differential gene expression between human and mouse microglia, which is enriched in Alzheimer’s and Parkinson’s disease genes, may be related to longevity. In this context, it is of note that cross-species evaluation of longevity-associated genes identified JAK/STAT signaling, amongst many other pathways (40). Thus, we speculate that evolution of longevity and immune function may be interrelated and may thus be reflected in mechanisms of age-related diseases.

Irrespective of the evolutionary reason(s) for LRRK2 being more highly expressed in microglia in humans compared to mice, these results have implications for our ability to model human diseases in other species. It is notable that mouse models where LRRK2 mutations have been introduced generally reproduce the biochemical effects of gain of function on Rab phosphorylation (41) but show only modest evidence of neurodegeneration, at least at endogenous expression levels (42). We posit that exposure to chronic inflammation may increase LRRK2 expression and activity in species that are genomically encoded to do so. Of note, the diminished age-dependent penetrance of LRRK2 mutations in humans, estimated to be below 30% of all carriers (43), may be related to non-genetically defined exposure. We speculate that immune activation, especially repeated or chronic febrile events, may increase PD risk, consistent with epidemiological data across neurodegenerative diseases (44). However, we caution against interpreting these data as suggesting that immune system function is a single causal determinant of PD risk. Recent data suggest that neuromelanin, which is found in vulnerable neuronal cells in humans but absent in many species used for laboratory modeling (45), may contribute to microglial activation and neuronal damage in the context of LRRK2 mutations (46).

Our data suggest that species differences may impact both CNS and peripheral immune cells in complex ways. For example, and consistent with recent data (24), we found that some murine monocytes did respond to IFN-ɣ with increased Lrrk2 expression, although this was quantitatively less than human monocytes. Thus, while our data show that human LRRK2 promoter elements are sufficient to infer enhanced responsiveness of murine microglia to IFN-ɣ, there are additional mechanisms that allow mouse cells to upregulate LRRK2 in the appropriate context. It is therefore of interest to note that peripheral exposure to LPS can induce LRRK2-dependent phenotypes where mouse peripheral cells are recruited to the CNS, albeit in the context of transgenic expression of a human BAC (15,16). It is also of note that mice expressing mutant Lrrk2 do show enhanced sensitivity to peripheral inflammatory phenotypes, including colitis that might secondarily lead to some loss of dopamine neurons in the substantia nigra (47,48). These observations demonstrate that understanding of cell- and signaling- contextual gene regulation is important for developing informative models of age-dependent chronic disease with direct human relevance.

The current study has several important limitations. While we have demonstrated that IFN-ɣ can increase LRRK2 expression in iPSC-derived cells and in human brain slices, the closest analogue of brain injection that we could access, we have not formally demonstrated that the same mechanisms as shown in iPSC cells are accessed *in vivo.* Equally, it is challenging to establish whether chronic inflammation contributes causally to diseases including PD and CD that have decades-long prodromal periods. Nonetheless, we expect that the current data will be helpful in selection of model systems and appropriate stimuli that might act as surrogates for human exposures. Our data presented here outline several novel mechanisms relevant to regulation of the PD and CD-associated gene LRRK2 that identify an unexpected species-dependent regulation of gene expression. These observations reinforce that genetic variations act in the context of physiological signaling pathways that are relevant for environmental exposures including common infections. We also note that there are likely mechanistic links between infection, inflammation and lysosome function such that maintenance of lysosomal membrane integrity may be required in acute circumstances but, speculatively, may be maladaptive over extended periods. Future work should incorporate this conceptual distinction between immediate responses to immune challenges and detrimental engagement of beneficial cellular responses in chronic conditions.

## Acknowledgements

We acknowledge the use of ChatGPT (developed by OpenAI) for code editing. All final content, including written and computational components, was reviewed and approved by the authors. We also thank Huikin Wang, National Institute for Mental Health, for assistance with preparing the human brain slices.

## Funding

This research was supported by the Intramural Research Program of the NIH, National institute on Aging and the National Institutes of Neurological Disorders and Stroke.

## Author contributions

AB and MRC conceptualized the primary experiments. AB, JP and NL conducted the experiments, supported by XR, HB, CK, AK, and JB. KZ and EM performed the human tissue extractions. Computational analyses were performed by AB, RC, JD, DA, JRG, AD and MRC. AB and MRC led the writing of this manuscript with contributions from all authors.

## Competing interests

The authors declare that they have no competing interests.

## Data and materials availability

All the data needed to evaluate the conclusions in the paper are present in the paper and/or the Supplementary Materials. Requests for resources and reagents should be directed to and will be fulfilled by the lead contact, Mark R Cookson. Reagents and methods in this study will be made available by the lead contact upon request.

Code is available at: https://github.com/neurogenetics/ADRD_iPSC

Raw data is available at: Multiome from human iPSC (reviewer token cbavueamjnezvct): https://www.ncbi.nlm.nih.gov/geo/query/acc.cgi?acc=GSE309354 ChipSeq (reviewer token ybsdmmqstlwdpsl): https://www.ncbi.nlm.nih.gov/geo/query/acc.cgi?acc=GSE309352 Mouse brain sn-RNAseq (reviewer token):

Human brain slice sn-RNAseq (reviewer token uhqnososfvwnpaz): https://www.ncbi.nlm.nih.gov/geo/query/acc.cgi?acc=GSE309355

## Materials and methods

### iPSC culture

All iPSC lines were expanded in Essential 8™ medium (Thermo scientific) on Matrigel coated dishes. Rock inhibitor was added to the medium for one day after thawing and after cell splitting. iPSCs were frozen using Synth-a-Freeze™ Cryopreservation Medium (Thermo scientific). iPS cells were passaged using TrypLE™ Select Enzyme (Thermo Scientific) or Cell dissociation Buffer (Thermo Scientific) for a total of 1-2 passages, and recovered post-split using Essential 8™ Medium with 10μM Rock inhibitor.

### Genome editing of A18945 iPSC line using RNPs

Alt-R CRISPR-Cas9 crRNAs were designed using the IDT CRISPR Custom Design Tool based on predicted high on-target activity and low off-target risk (see Table for sequences). crRNA and tracrRNA (IDT) were each resuspended to 200 µM in duplex buffer. To form the guide duplex, 5 µL crRNA and 5 µL tracrRNA were mixed, heated at 95 °C for 5 min, and cooled to room temperature. RNP complexes were prepared by combining 1.7 µL Cas9 nuclease (104 pmol), 1.2 µL crRNA:tracrRNA duplex (120 pmol), and 2.1 µL 1× PBS, followed by a 30-min incubation at room temperature. iPSCs at 70–80% confluence were dissociated with Accutase and counted. For each reaction, 8×10⁵ cells were pelleted (1000 rpm, 3 min) and resuspended in 100 µL P3 Primary Cell Solution (Lonza). Five microliters of pre-assembled RNP were added to the cells, mixed gently, and transferred to a 100 µL nucleocuvette. Cells were nucleofected using the Primary Cell P3 program with CA-137 pulse code. Following nucleofection, cells were transferred to Matrigel-coated 6-well plates containing E8 medium with 10 µM ROCK inhibitor and cultured at 32 °C/5% CO₂ for 2 days before returning to 37 °C. Edited pools were expanded and frozen. Single-cell-derived clones were screened for peak A and peak B deletions using primers listed in the Table. Clones with deletions on both alleles were validated by sequencing. The CRISPR/Cas9-induced homozygous deletion in the ΔA/ΔA clone spans chr12: [40223572]–[40223984] (hg38). The ΔB/ΔB clone carries a compound heterozygous deletion, chr12:40220114–40220646 on one allele and chr12:40220124–40220647 on the second allele (Figure S2).

### iMicroglia Differentiation

iPSCs were cultured in E8 media without ROCK for at least 2 days before beginning the differentiation as described (26). On the first day of the differentiation, 10,000 cells were plated per well in 96-well V-bottom shapes ultra-low attachment plates (Sunoko) in Embryoid body medium (EBM) containing E8 media, 10 mM ROCK inhibitor, 50 ng/ml BMP-4, 20 ng/mL SCF,and 50 ng/mL VEGF-121. Next day, 80 μM of media was replaced with 100 μM of EBM without ROCK inhibitor. Culture EBs for additional 2 days, with EBM medium change every day. On day 4, 20-24 EBs were transferred to 1 well of a 6-well plate and 3 mL of Microglia progenitor media (MPM) containing X-VIVO 15 Lonza media, 2 mM GlutaMax, 55 mM beta-Mercaptoethanol, 100 ng/mL M-CSF and 25 ng/mL IL-3 was added. After adherence of EBs in a non-coated cell culture dish for 3-4 days, media was changed every other day for fresh MPM. Large amounts of microglia progenitor cells usually appeared on day 12-day 18. Microglia progenitor cells were collected by centrifugation and 0.6-0.7 million cells per 1well of the non-coated 6-well plate were seeded for the final differentiation in Microglia maturation media (MMM) containing Advanced RPMI, 2 mM GlutaMax, 100 ng/mL IL-34 and 10 ng/mL GM-CSF. Cells were further differentiated for additional 10-12 days with full MMM media changes every other day.

### Neuronal Differentiation

iPSCs were differentiated to forebrain neurons (FBn) as previously described (49) with some modifications. iPSC lines were plated in matrigel coated 6 well plates and grown to 90% confluence. Upon reaching 90% confluence, FBn N3 differentiation media (50% DMEM/F12 (Thermo Fisher, 11320033), 50% Neurobasal (Thermo Fisher, 21103049) with 0.5x GlutaMAX (Thermo Fisher, 25030-081), 1x Penicillin-Streptomycin (Thermo Fisher, 15140122), 0.5x B-27 minus vitamin A (Thermo Fisher, 12587010), 0.5x N2 supplement (Thermo Fisher, 17502048), 0.5x MEM Non-Essential Amino Acids (NEAA) (Thermo Fisher, 11140-050), 0.055 mM 2-mercaptoethanol (Thermo Fisher, 21985-023) and 1 µg/mL Insulin (Millipore Sigma, 91077C)) plus 1.5 µM Dorsomorphin (Tocris Bioscience, 3093) and 10 µM SB431542 (Stemgent, 04-0010-05) was added and changed daily until differentiation day 11. N3 without Dorsomorphin and SB431542 was replaced daily until day 16. From day 16 to 19, N3 media with 0.05uM Retinoic acid (Sigma, R2625) was changed every day. Wells were split 1:2 on day 19 using accutase (Sigma-Aldrich, A6964). Cells were replated in N3+RA+ ROCK inhibitor/ Y-27632 (Stemcell Technologies, 72302) on Poly-L-Ornithine (Sigma, 27378-049), Fibronectin (Fisher Scientific, CB40008A), Laminin (Sigma, L6274-.5MG) and matrigel coated plates. Media was changed to N4 the following day and every other day after until differentiation day 60. DAN differentiation was performed as described previously (50).

### Single Cell Multiome RNA and ATAC sequencing

Differentiated iPSc-derived microglia cells were washed with 1x PBS, Accutase solution was added to each well and cells were incubated at RT for 10 min. Next, cells were collected in 1xPBS by centrifugation, 300g, 5min and washed once with 1xPBS + 0.04% BSA. 100 μl of chilled Lysis Buffer (10mM Tris-HCl (pH 7.4), 10mM NaCl, 3mM MgCl, 1% BSA, 0.1% Tween-20, 0.1% NP40, 0.01% Digitonin) were added to 500,000 cells, cells were resuspended 10 times, and incubated on ice for 5 min. Next, 1 ml of chilled Wash Buffer (10mM Tris-HCl (pH 7.4), 10mM NaCl, 3mM MgCl, 1% BSA, 0.1% Tween-20) was added to the lysed cells and mixture was pipetted 5 times. Nuclei were precipitated at 500g for 5 min at 4°C. Nuclei were washed in the Wash Buffer for a total of 3 washes. 20, 000 nuclei/sample were processed according to Chromium next GEM Single cell Multiome ATAC+Gene Expression user guide (CG000338 Rev E). Chromium Single Cell Multiome Gene Expression and ATAC libraries were sequenced using standard Illumina® paired-end sequencing with P5 and P7 primers. Library concentration and sizes were determined using the Agilent 2100 Bioanalyzer with the High Sensitivity DNA kit (Agilent, 5067-4626). Paired end sequencing was completed as recommended on a NovaSeq6000 with 28,91 paired end reads for single cell RNA-seq and 50, 49 paired end reads for single cell ATAC-seq samples. NICS did sample processing using the count method from CellRanger (v5.0.1, introns included and refdata-gex-GRCh38-2020-A annotation file) and CellRanger-atac (v2.0.0, refdata-cellranger-arc-GRCh38-2020-A-2.0.0 annotation file) tool packages for RNA and ATAC respectively. Single-cell multiome libraries were sequenced to a mean depth ranging from 15,413 to 32,278 raw read pairs per cell for chromatin accessibility and 15,520 to 35,149 raw read pairs per cell for gene expression across samples. Sequencing depth was comparable across experimental conditions and within recommended ranges for the 10x Genomics Single Cell Multiome ATAC + Gene Expression platform. For multiome data, low-quality cells were removed based on chromatin accessibility and gene expression metrics. Cells with fewer than 1,000 or more than 100,000 ATAC fragments, nucleosome signal ≥2, or low TSS enrichment were excluded. For gene expression, cells with fewer than 200 or more than 25,000 RNA counts were removed. Mitochondrial read percentage was calculated and regressed out during SCTransform normalization.

### siRNA screen

Dharmacon on-target plus, smart pool cherry pick siRNA library was made targeting 171 genes. Non-targeting control siRNA was included in each plate. siRNA was diluted to 2uM concentration using 1x siRNA dilution buffer (Dharmacon). Fully differentiated microglia were plated 20x10^3^ cells/well, in 96 well plates. Next day microglia were transfected with 5nM siRNA using Glia Mag transfection reagent (Oz Biosciences) on magnetic plates for 30 min. Magnetofection was repeated again the next day for a total of two days. 48 hours after siRNA transfection microglia were treated with 20 ng/ul IFN-ɣ for 24 hours and plates were processed using TaqMan Fast Advanced Cells-to-Ct Kit (Thermo Scientific, cat # A35377). 24h after treatment cells were first washed in 1x PBS and then 50 μL of Cells-to-CT Lysis Solution (Thermo Scientific, cat # A35377) were added to each well and cells were incubated at RT for 5 min on orbital shakers. Prepared lysates were used for RT first and then for qPCR according to TaqMan Fast Advanced Cells-to-CT protocol (Thermo Scientific, cat # A35377). LRRK2 mRNA levels were measured using FAM-MGB probes (Thermo Fisher Scientific, cat #4331182) VIC-MGB probe for control PPID gene was included in each reaction (Thermo Fisher Scientific, Hs00234593_m1). LRRK2 expression value was normalized to PPID for each sample. For each siRNA condition, statistical comparisons were performed against the non-targeting control (NTC) using two-sided t-tests. Corresponding fold changes were calculated, and false discovery rate (FDR) correction was applied to the resulting p-values to account for multiple testing.

### Chromatin immunoprecipitation (ChIP)

Chromatin immunoprecipitation (ChIP) was performed using the SimpleChIP® Plus Enzymatic Chromatin IP Kit (Magnetic Beads) (Cell Signaling Technology, Cat# 9005, Danvers, MA, USA) following the manufacturer’s protocol, with slight modifications. Briefly, iPSC-derived microglia cells (iMGL) were seeded in 10 cm culture dish at a concentration of 4 × 10^6^ cells/ml and crosslinked with 1% formaldehyde for 10 min at room temperature. The crosslinking reaction was stopped by adding 10X glycine for 5 min at room temperature (RT). After washing twice with cold PBS, cells were collected by scraping in PBS supplemented with Protease inhibitor cocktail (PIC). Nuclei were prepared, and chromatin was incubated with micrococcal nuclease at 37 °C for 20 min, which was followed by 30 sec with sonication to digest DNA to the optimal length (150-900 bp). The supernatants were immunoprecipitated by being incubated with normal IgG (Cell Signaling Technology, Cat # 2729), Histone H3 (D2B12) (Cell Signaling Technology, Cat# 4620), and anti-STAT1(Cell Signaling Technology, Cat # 9172) at 4 °C for overnight. The next day, the immunocomplexes were rotationally incubated with 30 µl of ChIP-Grade Protein G Magnetic Beads (Cell Signaling Technology, Cat # #9006) for 2 h at 4 °C and then were washed three times using low salt wash buffer and 1 time with high salt wash buffer at 4 °C for 5 min per wash. Bead-bound chromatin was eluted by ChIP elution buffer for 30 min at 65 °C with gentle vortex mixing (1200 rpm) and crosslinks were reversed by treatment with 5 M NaCl and proteinase K for 2 h at 65 °C. ChIP DNA was purified using spin columns and subsequently quantified by quantitative real-time PCR (qPCR). ChIP DNA libraries were prepared using the Illumina ChIP-Seq DNA Sample Prep Kit (Illumina,IP-102-1001). Final libraries were purified, quantified, and quality-checked before sequencing on an Illumina platform. ENCODE4 transcription factor ChIP-seq pipeline was utilized to map sequence reads to human reference GRCh38, identify signals, and peaks calling. The Irreproducible Discovery Rate (IDR) was used to evaluate consistency of ChIP-seq experiments with replicates and to create the optimal set of more sensitive peaks. The signal coverages at nucleotide resolution are reported as signal p-values and fold change over control at each position. bigWig, narrowPeak, as fold-over control at each position, and as a p-value to reject the null hypothesis that the signal at that location is present in the control (except for Input samples).

### siRNA transfection

iMicroglia were transfected with magnetic particles (OZ bioscience, Cat# KGL0250, Marseille, France) using targeting siRNA (STAT1, IRF1, JAK1, and JAK2) following the manufacturer’s instructions. Briefly, iMicroglia were seeded in 12-well plates at a concentration of 0.84 × 10^6^ cells/ml and cultured overnight, after which control siRNA, targeting siRNA, and glial-mag (GM) transfection reagent (OZ bioscience, Cat# KGL0250, Marseille, France) were added to these cells. Next, a glial-boost (GB) solution was added to promote transfection efficiency. Cell cultures were then placed on a magnetic plate, incubated for 30 min, afterwards the magnet removed, and cell culture media were added, and cells incubated at 37°C for a further 24 h period. The extent of knockdown was evaluated by western blotting analysis.

### SDS PAGE and Western Blotting

Proteins were resolved on 4–20% Criterion TGX pre-cast gels (Biorad) in SDS/Tris-glycine running buffer and transferred to membranes by semi-dry trans-Blot Turbo transfer system (Biorad). The membranes were blocked with Odyssey Blocking Buffer (Licor Cat #927-40000) and then incubated for 1h at RT or overnight at 4°C with the indicated primary antibody. The membranes were washed in TBST (3×5 min) at room temperature (RT) followed by incubation for 1h at RT with fluorescently conjugated goat anti-mouse or rabbit IRDye 680 or 800 antibodies (LICOR). The blots were washed in TBST (3×5 min) at RT and scanned on an ODYSSEY^®^ CLx (Licor). Quantitation of western blots was performed using Image Studio (Licor).

### Mouse Interferon-γ Injection

Mice were housed in a 12-h light/dark cycle with ad libitum access to food and water. All animal work was performed in accordance with the Institutional Animal Care and Use Committee (ACUC) of the National Institute on Aging (NIA/NIH). Strains C57Bl/6J (WT; Jax) and C57Bl/6J -Lrrk2tm1.1Cai/J (LRRK2 KO; Jax) were used for this experiment. Three-months-old mice (n=4 per group) were initially anesthetized by 5% iso-flurane and kept under anesthesia using 1–2% isoflurane. Mice were placed into a stereotaxic frame (Kopf) and the skull was exposed by making an incision down the midline. A hole was drilled into the skull at 4 injection sites. Coordinates were as follows: anteroposterior +0.2 mm, mediolateral ±2.0 mm from bregma for striatum (bilateral); and anteroposterior −3.3 mm, mediolateral ±1.5 mm for substantia nigra (bilateral). A pulled glass capillary (blunt) attached to a 5 μl Hamilton syringe was used for injection. First, an air bubble of 1 μl was pulled in followed by 1 μl of 20 ng/μl of mouse INFγ solution (20 ng total) or PBS as control per injection site. The capillary was lowered to dorsoventral −3.2 mm from bregma for the dorsal striatum and −4.3 mm for substantia nigra. The solution was delivered at a rate of 0.4 μl/min. After the injection, the capillary was held in place for 2 min, and then slowly withdrawn from the brain. The wound was closed using clips and mice recovered on a heating pad with ketoprofen (5mg/kg) as analgesic. After 24hr, mice were euthanized using CO2, brains were taken out and washed in ice cold PBS. The striatum (average 15.0 mg ± 3.3) as well as the ventral midbrain (average 15.6 mg ± 2.6) were dissected, flash frozen and stored at −80 until processed for Single Cell Multiome or western blot. For immunohistochemistry, animals were deeply anesthetized using an overdose of sodium pentobarbital (30 mg/kg). Mice were perfused via the ascending aorta first with 10 ml of 0.9% NaCl followed by 50 ml of ice-cold 4% paraformaldehyde (PFA in 0.1 mM phosphate buffer, pH 7.4) for 5 min. Brains were removed and post-fixed in 4% PFA for 24 h and then transferred into30% sucrose for cryoprotection. The brains were then cut into 30 um thick coronal sections (microtome Leica) and stored in an antifreeze solution (0.5 mM phosphate buffer, 30% glycerol, 30% ethylene glycol) at –20◦C until further processed. After 24 hours, striatum and substantia nigra were collected for nuclei purification using 10x Genomics single-cell nuclei isolation kit (PN-1000494). 20, 000 nuclei/sample were processed according to Chromium next GEM Single cell Multiome ATAC+Gene Expression user guide (CG000338 Rev E). Chromium Single Cell Multiome Gene Expression and ATAC libraries were sequenced using standard Illumina® paired-end sequencing with P5 and P7 primers. Paired-end sequencing was performed on a NovaSeq 6000. Samples were processed using the cellranger count pipeline (v7.1.0) for gene expression data, including intronic reads, with the reference genome refdata-gex-mm10-2020-A. For ATAC-seq data, the cellranger-atac count pipeline (v2.0.0) was used with the refdata-cellranger-arc-mm10-2020-A-2.0.0 reference package. Single-cell RNA-seq libraries from substantia nigra samples were sequenced to a mean depth of 20,453 ± 4,072 raw reads per cell (mean ± SD), while chromatin accessibility libraries were sequenced to a mean depth of 33,659 ± 12,372 raw reads per cell.Single-cell RNA-seq libraries from striatum samples were sequenced to a mean depth of 29,684 ± 3,125 raw reads per cell (mean ± SD), with chromatin accessibility libraries sequenced to a mean depth of 32,442 ± 7,083 raw reads per cell. Sequencing depth was comparable across experimental conditions and within recommended ranges for the 10x Genomics Single Cell Multiome ATAC + Gene Expression platform. For multiome data, low-quality cells were removed based on chromatin accessibility and gene expression metrics. Cells with fewer than 1,000 or more than 100,000 ATAC fragments, nucleosome signal ≥2, or low TSS enrichment were excluded. For gene expression, cells with fewer than 200 or more than 25,000 RNA counts were removed. Mitochondrial read percentage was calculated and regressed out during SCTransform normalization.

### Generation of Monocytes, Macrophages, and Dendritic Cells from BMDCs

Human Bone Marrow Mononuclear Cells (BM-MNCs) derived from three human different donors were purchased from StemCell Technologies (Catalog #70001). BMDCs were differentiated into human or mouse monocytes by culturing the BM-MNCs for 4 days in RPMI 1640 medium supplemented with 10% FBS and either human or mouse M-CSF (20 ng/mL). Monocytes were then either harvested for Western blot analysis or further differentiated into macrophages or dendritic cells. For macrophage differentiation, monocytes were cultured in the presence of M-CSF (20 ng/mL) for an additional 4 days. For dendritic cell differentiation, monocytes were cultured with GM-CSF (20 ng/mL) and IL-4 (20 ng/mL) for additional 4 days, with a media change on day 3. All cells were maintained at 37°C in a humidified atmosphere with 5% CO₂, and media were refreshed every 2–3 days unless otherwise specified. Monocytes, macrophages, and dendritic cells were treated with 20 ng/µL human or mouse IFN-γ for 24 or 48 hours.

### Cellular Treatments

iPSC-derived microglia, forebrain neurons (FBNs), dopaminergic neurons (DANs), and human bone-marrow–derived monocytes, macrophages, and dendritic cells, as well as human brain slice cultures, were treated with 20 ng/mL ( ≥ 200 IU/mL) human IFN-γ (PeproTech, Cat. #300-02-100 UG) or 20 ng/mL (≥ 100 IU/mL) mouse IFN-γ (PeproTech, Cat. #315-05-20UG) for the designated time points. Human-derived cultures were treated with human IFN-γ, whereas mouse-derived cultures were treated with mouse IFN-γ, as human IFN-γ is not active in mouse cells.

For in vivo experiments, 1 µL of mouse IFN-γ solution (20 ng/mL; ≥ 1 × 10⁵ IU/mL; 20 ng total) or PBS (control) was injected per injection site.

iPSC-derived microglia were treated with 20 ng/mL recombinant human IL-4 (PeproTech, Cat. #200-04; ≥ 100 IU/mL), recombinant human IL-10 (PeproTech, Cat. #200-10; ≥ 100 IU/mL), recombinant human IFN-β (PeproTech, Cat. #300-02BC; ≥ 200 IU/mL), or recombinant human IFN-λ1 (IL-29) (PeproTech, Cat. #300-02L; ≥ 400 IU/mL) for 24 hours.

iPSC-derived microglia were also treated with 1 mM LLOME for 15 minutes.

### Human Brain Slice Culture and Single-Nucleus RNA Sequencing

Tissue blocks (5–7 mm) of human cortical brain tissue obtained from epilepsy surgery patients were sectioned in the lab using a vibratome at ∼7–11 °C to preserve ultrastructure. One section per individual was used in this study. We obtained human brain sections from two individuals: frontal lobe tissue from a 27-year-old female and anterior temporal lobe cortex from a 36-year-old male. Three hundred μm sections were evenly cut in half and incubated in artificial cerebrospinal fluid (126 mM NaCl, 2.5 mM KCl, 1.25 mM NaH₂PO₄, 26 mM NaHCO₃, 2 mM CaCl₂, 1–2 mM MgCl₂, and 10 mM glucose; pH ∼7.3–7.4) in the presence or absence of 25 ng/mL IFN-γ for 6 hours. After incubation, sections were frozen and processed using the 10x Genomics single-cell nuclei isolation kit (PN-1000494), following the manufacturer’s protocol. Approximately 10,000 nuclei per sample per treatment were isolated and processed using the 10x Genomics Single Cell 3’ Gene Expression kit (cat #PN-1000121). Paired end sequencing was completed on an Illumina NovaSeq X Plus. Samples were processed using the count method from CellRanger (v7.2.0, introns included and refdata-gex-GRCh38-2024-A annotation file) tool packages for RNA. Human brain single-cell RNA-seq libraries were sequenced to a mean depth ranging from 48,642 to 61,459 raw reads per cell across individuals and treatment conditions. For gene expression, cells with fewer than 200 or more than 25,000 RNA counts were removed. Mitochondrial read percentage was calculated and regressed out during SCTransform normalization.

### Processing of single cell datasets

Single cell or single nuclei RNA or ATAC data were analyzed using the R package Seurat. Full code is available (github). Key marker genes for mouse substantia nigra and striatum were taken from prior literature (51).

### Comparative genomics

Peak locations (in hg38) were used as input into the UCSC Genome Browser (52). For these loci, we obtained multiple species alignments from the “Cactus 447-way” vertebrate alignment track. Alignments were produced using the Cactus algorithm, which employs a progressive, graph-based approach to align vertebrate genomes while handling structural variations and paralogy through cactus graphs (53). The alignment pipeline began with the Zoonomia 241 species alignment (54,55) as a starting point, removed all primates and outdated assemblies, then incorporated alignments from 233 newly sequenced primate genomes(56,57).

For visualization, sequences were compared to the human reference (hg38), where matches represent identical nucleotides, mismatches represent divergent nucleotides, and gaps represent insertions/deletions or assembly gaps in non-human species. Multiple alignment blocks were concatenated with hg38 serving as the gap-free reference sequence. A time-calibrated phylogenetic tree for select mammalian species was obtained from TimeTree (58), which provides divergence times based on molecular clock analyses and fossil calibrations across vertebrate taxa.

Selected species included *Pan troglodytes* (chimpanzee), *Gorilla gorilla* (western gorilla), *Pongo abelii* (Sumatran orangutan), *Macaca nemestrina* (pig-tailed macaque), *Macaca mulatta* (rhesus macaque), *Cercocebus atys* (sooty mangabey), *Papio anubis* (olive baboon), *Alouatta palliata* (mantled howler monkey), *Ateles geoffroyi* (Geoffroy’s spider monkey), *Cebus albifrons* (white-fronted capuchin), *Callithrix jacchus* (common marmoset), *Aotus nancymaae* (Ma’s night monkey), *Eulemur flavifrons* (blue-eyed black lemur), *Lemur catta* (ring-tailed lemur), *Microcebus murinus* (gray mouse lemur), *Daubentonia madagascariensis* (aye-aye), *Galeopterus variegatus* (Sunda flying lemur), *Mus spretus* (Algerian mouse), *Mus musculus* (house mouse), *Rattus norvegicus* (Norway rat), *Cricetulus griseus* (Chinese hamster), *Mesocricetus auratus* (golden hamster), *Sus scrofa* (wild boar), *Ovis canadensis* (bighorn sheep), *Hippopotamus amphibius* (hippopotamus), *Mesoplodon bidens* (Sowerby’s beaked whale), *Equus caballus* (horse), *Zalophus californianus* (California sea lion), *Felis catus* (domestic cat), *Canis lupus familiaris* (domestic dog), and *Tolypeutes matacus* (southern three-banded armadillo). Position Weight Matrices (PWMs) for transcription factor binding sites were obtained from the HOCOMOCO v11 database of human transcription factor binding models(59). Transcription factor binding sites in the peak sequences (hg38) were identified using the *searchSeq* function in the R package *TFBSTools* (60) with a minimum score threshold of 70%. We visualized the locations of the resulting STAT1 binding sites.

## STAR*METHODS

### KEY RESOURCES TABLE

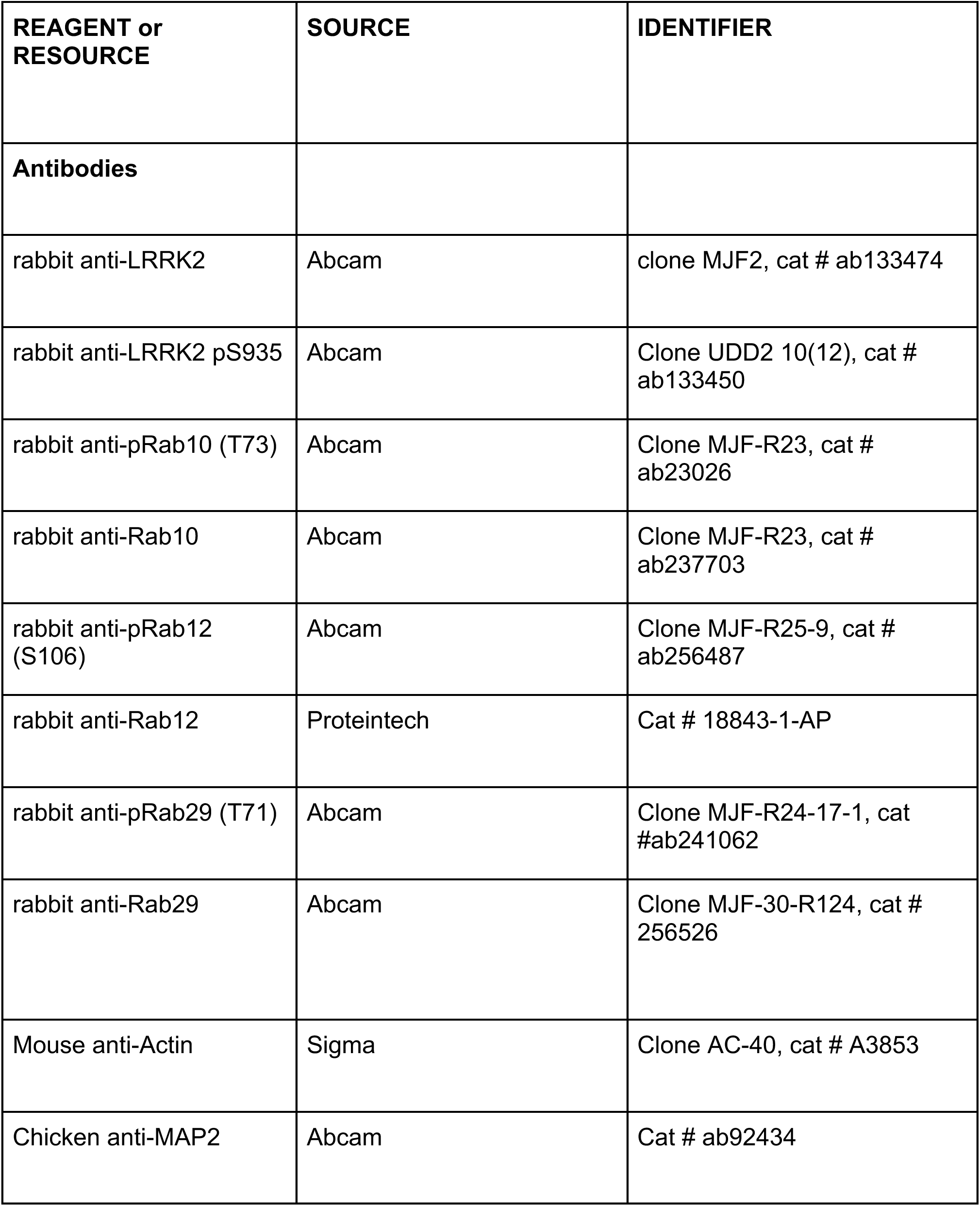
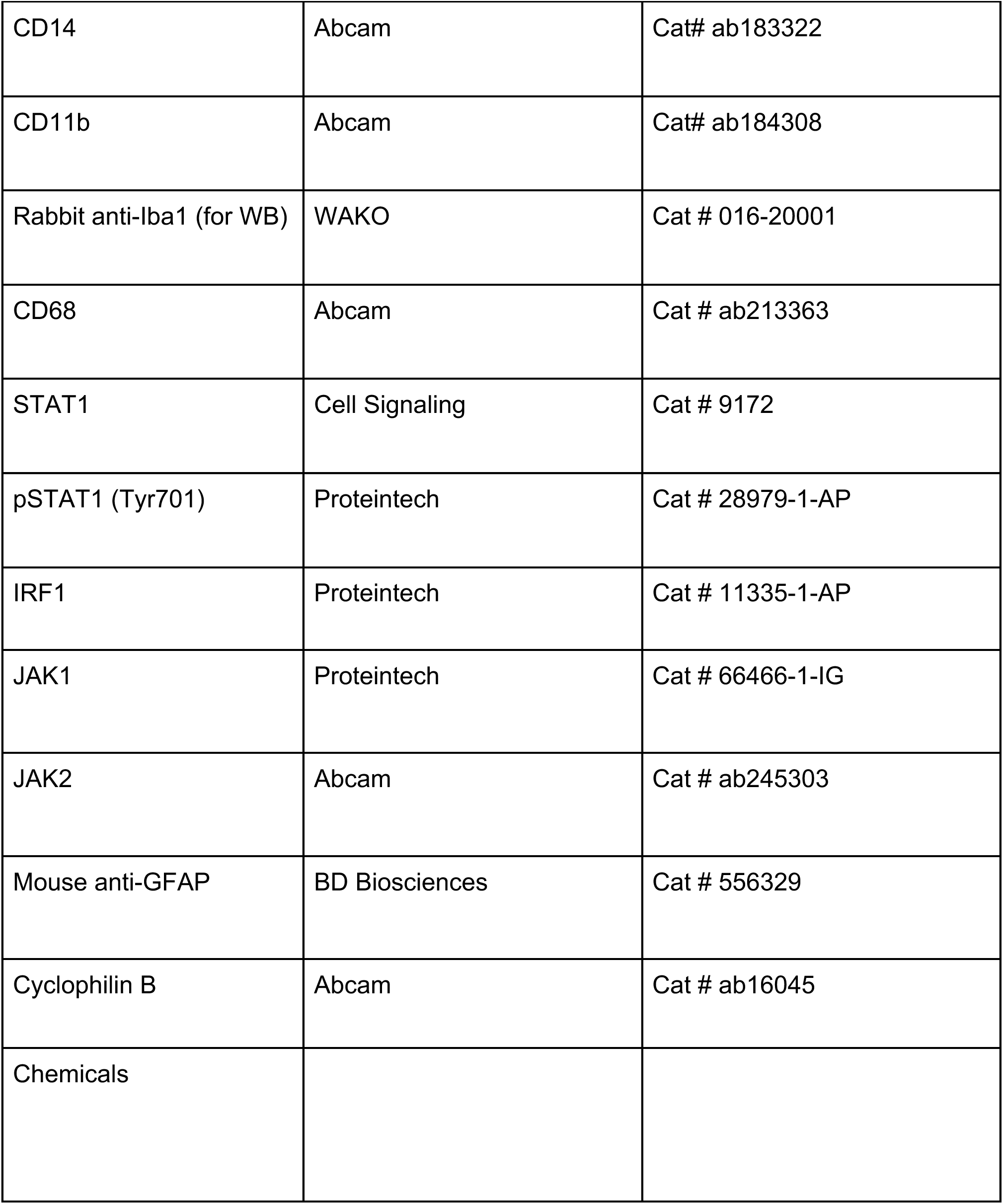
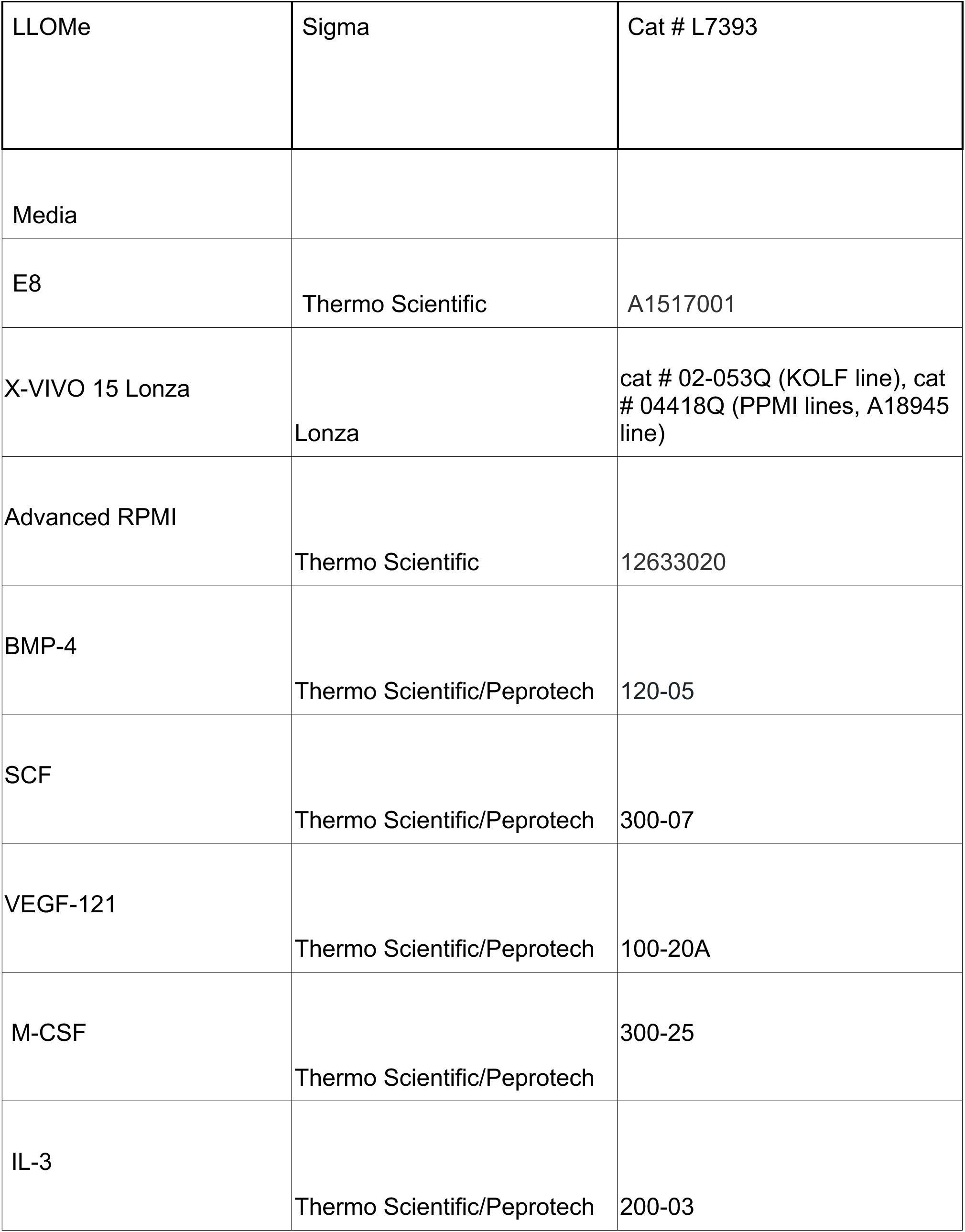
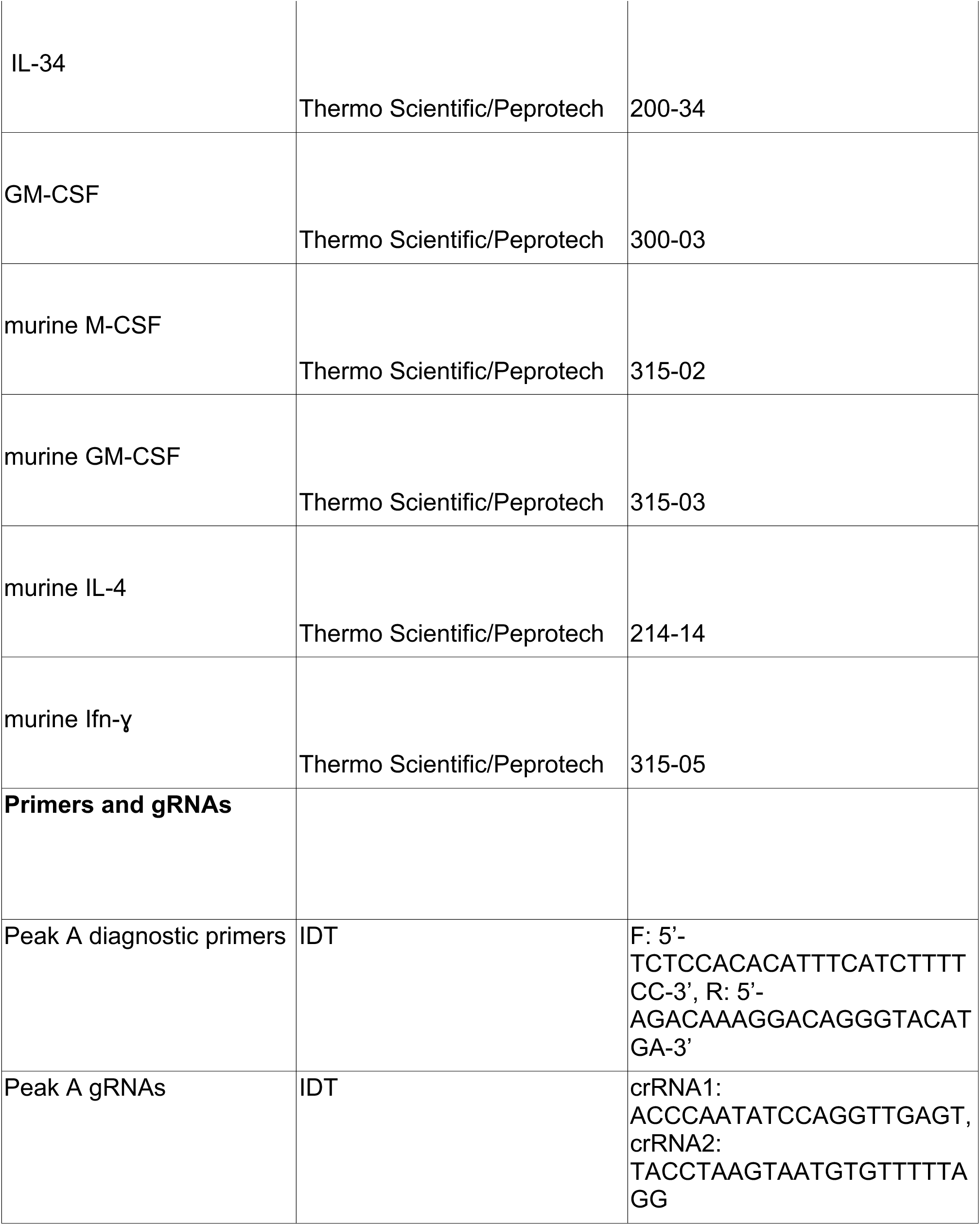
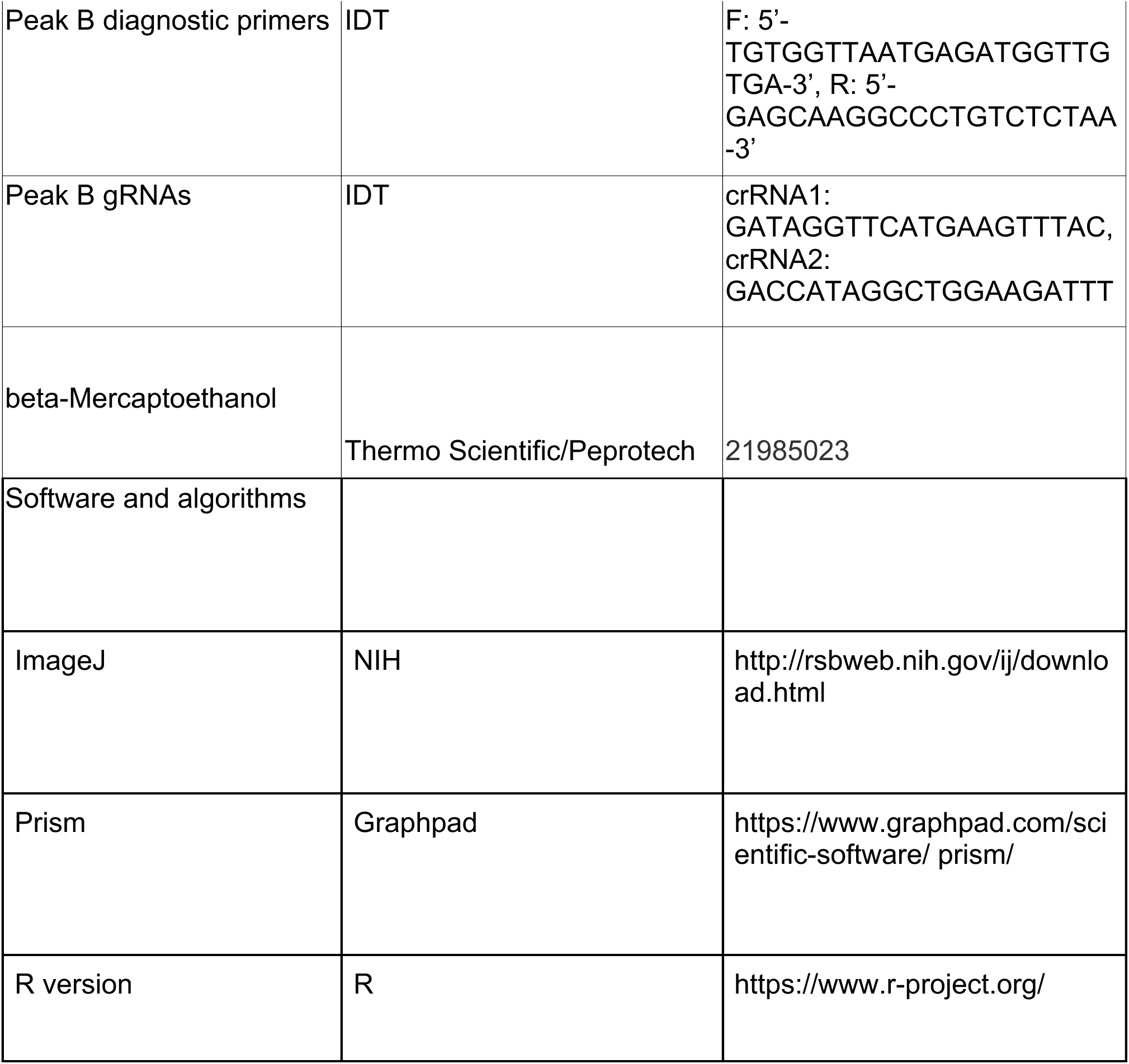

## Supplementary figure legends

**Figure S1.**
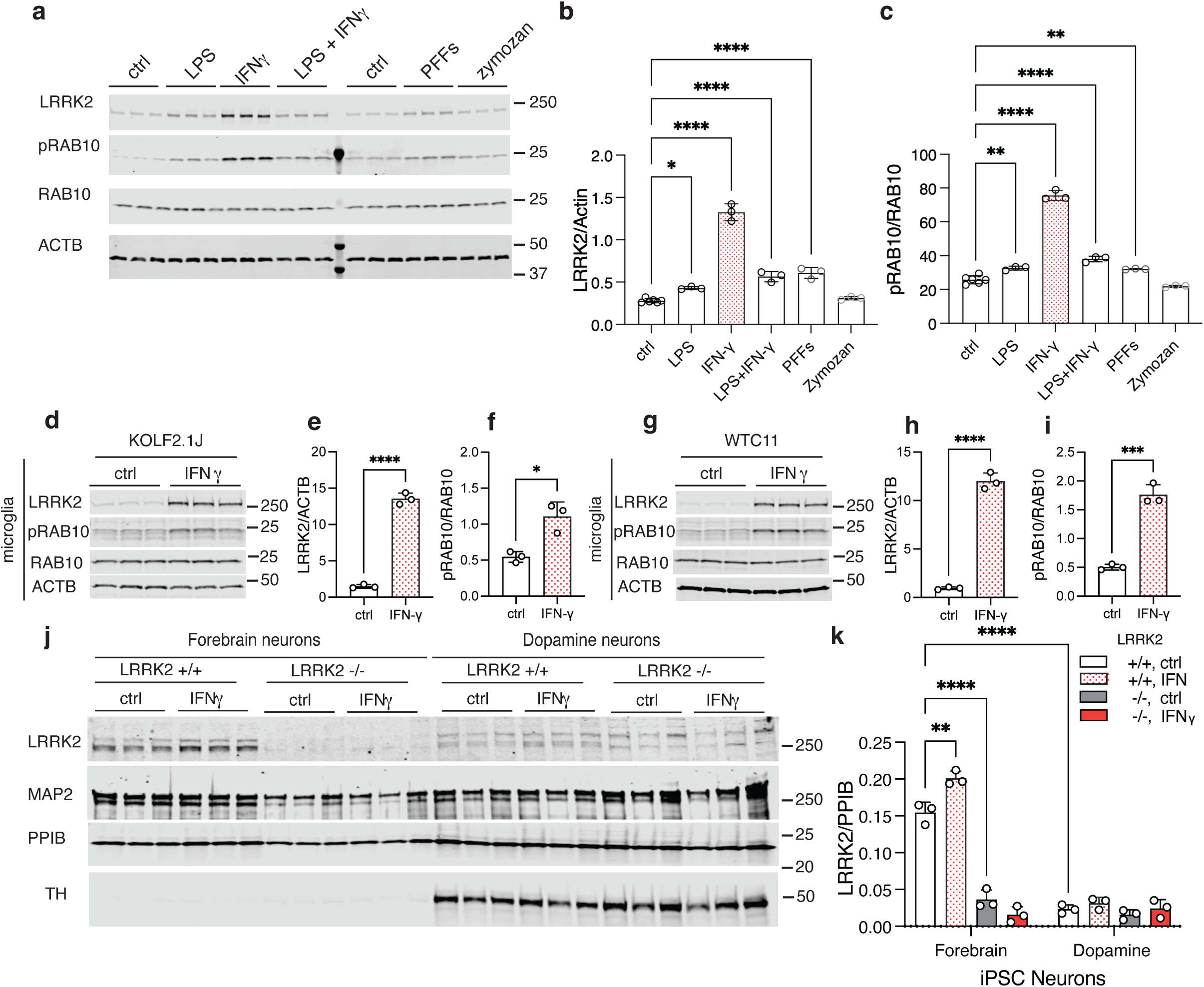
Induction of LRRK2 by inflammatory stimuli across iPSC-derived cells. (a) LRRK2 expression and activity was assessed in iMicroglia following stimulation with LPS, IFN-ɣ, a combination of LPS and IFN-ɣ, α-synuclein preformed fibrils (PFFs), or zymosan. Among these, IFN-ɣ elicited the strongest induction of LRRK2. However, both LPS and PFFs also led to significant increases in LRRK2 expression (b) and Rab10 phosphorylation (c). *, *p*<0.05; ** *p* <0.001; ***, *p*<0.0001, ****, *p*<0.0001 by one-way ANOVA for treatment (LRRK2, F_5,15_ = 184.8; pRAB10, F_5,15_ = 334.1) with Tukey’s *post-hoc* test to compare all conditions. For clarity, only comparisons to control conditions (ctrl) are shown. (d-g) LRRK2 induction was evident in iMicroglia derived from other iPSC lines, KOLF2.1J (d, quantified for LRRK2 in e; ****, *p*<0.0001 by two-tailed t-test; t=24.9, df=4; quantified for pRAB10 in f; *, *p*=0.0107 by two-tailed t-test; t=4.518, df=4, n=9, three replicate cultures across three independent differentiations), WTC11 (g, quantified for LRRK2 in h; ****, *p*<0.0001 by two-tailed t-test; t=22.09, df=4; quantified for pRAB10 in i; ***, *p* =0.0003 by two-tailed t-test; t=11.68, df=4, n=9, three replicate cultures across three independent differentiations). LRRK2 induction was also observed in 35-day-old forebrain neurons (j, quantified in k; **, *p*<0.01 by two-tailed t-test; t=8.57, df=4, n=3 replicate cultures), but not in dopaminergic neurons (j, quantified in k; ns, *p*=0.5816 by two-tailed t-test; t=0.5987, df=4, n=3 replicate cultures) differentiated from the A18945 iPSC line. For all blots, each lane is a replicate culture and molecular weight markers are shown on the right. For all graphs, bars show mean values, error bars are standard deviation and individual points represent independent cultures.

**Figure S2.**
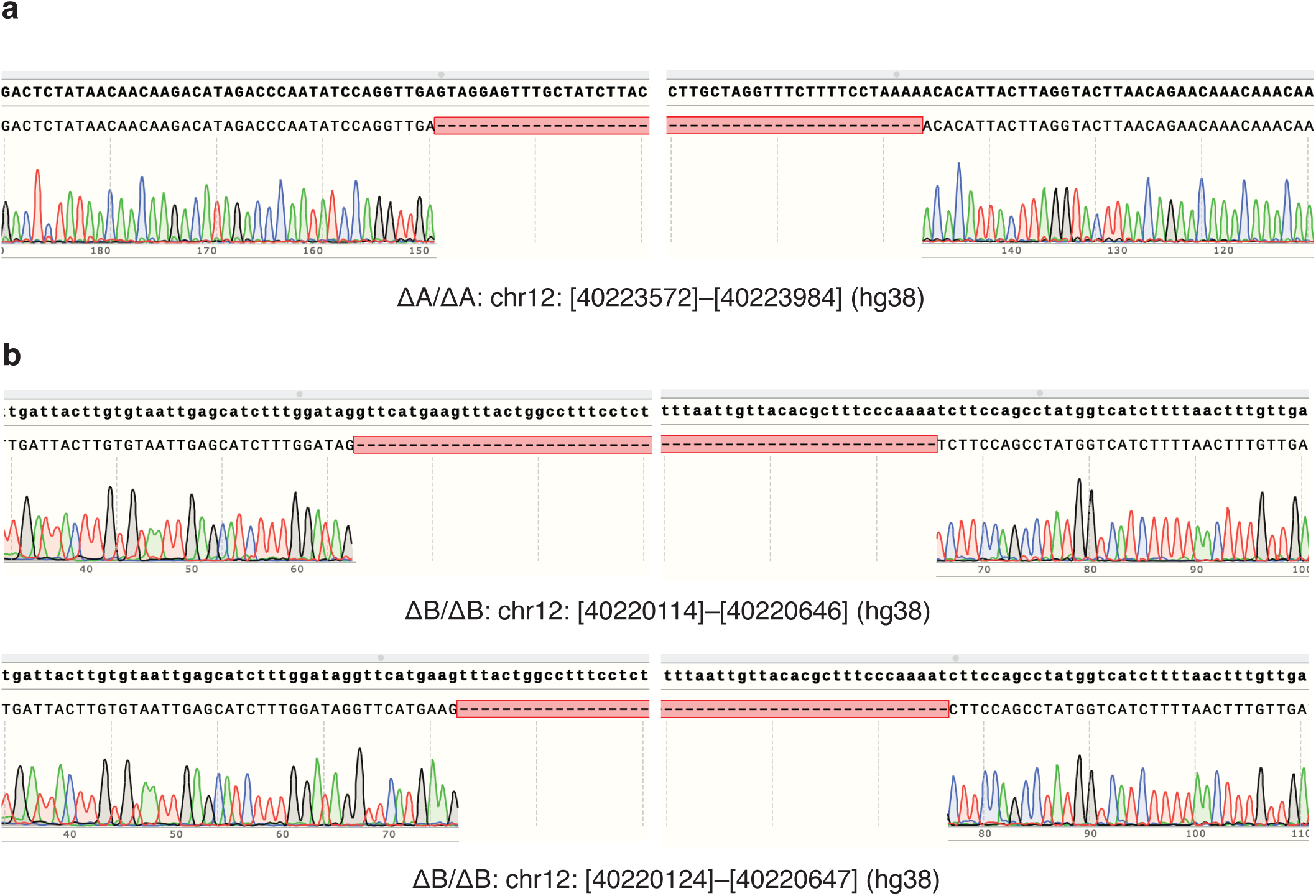
Sequence confirmation of CRISPR deletion clones. (a) Sanger sequencing of the ΔA/ΔA clone revealed a homozygous 412 bp deletion spanning chr12:40223572–40223984 (hg38). (b) Sanger sequencing of the ΔB/ΔB clone revealed a compound heterozygous deletion, with a 532 bp deletion spanning chr12:40220114–40220646 on one allele and a 523 bp deletion spanning chr12:40220124–40220647 on the second allele.

**Figure S3.**
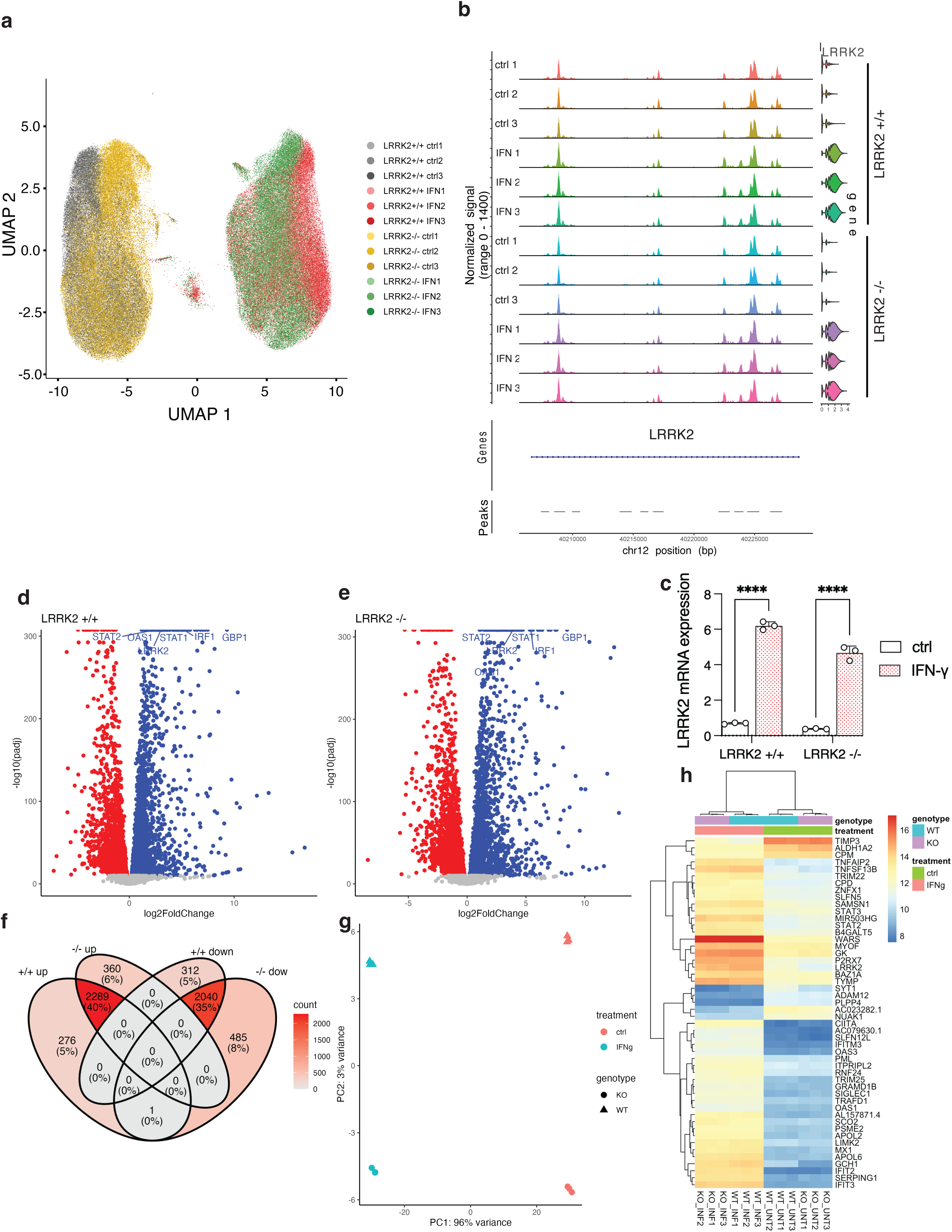
Changes in chromatin accessibility and gene expression after IFN-ɣ treatment. (a) UMAP feature plots of iMicroglia derived from LRRK2 +/+ and LRRK2 -/- iPSC lines colored by independent cultures, showing separation of treatment by the first UMAP dimension. A total of 196,320 **cells** are shown. (b) ATAC reads in LRRK2 +/+ and LRRK2 -/- samples (n=3 independent cultures per genotype and treatment) without (ctrl) or with IFN-ɣ treatment, shown alongside a violin plot displaying the distribution of LRRK2 RNA expression in the same cells (right). Appearance of Peak A (ch12:40223500-40224000) is evident in all samples treated with IFN-ɣ but not controls. The intensities of Peak B (ch12:40220110-40220600) and Peak C (ch12:40209200-40209000) were increased following IFN-ɣ stimulation. (c) Quantification of LRRK2 mRNA expression levels in multiome samples derived from WT and LRRK2 KO iMicroglia. *** *p* <0.0001; ****, *p*<0.00001 by two-way ANOVA for treatment by genotype (F_1,8_ treatment = 1384; F_1,8_ genotype = 48.74; F_1,8_ interaction = 21.59, n=3 independent cultures per genotype and treatment). For clarity, only significant comparisons are indicated: the effect of genotype in untreated cells was not significant (p=0.4056). (d, e) Volcano plots from pseudobulked expression data for LRRK2 +/+ iMicroglia (d) and LRRK2 -/- cells (e) in response to IFN-ɣ treatment. Significantly differentially expressed genes log2 fold change greater > 2 and an adjusted p < 0.05 are indicated in blue for upregulated and red for downregulated genes. (f) Venn diagram showing that a total of 2289 genes were upregulated and 2040 were downregulated concordantly in both genotypes, with no genes exhibiting discordant directions of change. (g) Principal component analysis (PCA) of the 500 most variable genes revealed that treatment was primarily associated with PC1, accounting for approximately 96% of the total variance, while genotype was linked to PC2, which explained only about 3% of the variance. (h) Heatmap of differentially expressed genes across experimental conditions. Each row represents a gene, and each column represents a sample. Gene expression values were normalized and scaled, with red indicating higher expression and blue indicating lower expression levels. Hierarchical clustering was performed on genes and samples based on similarity in expression profiles using Euclidean distance.

**Figure S4.**
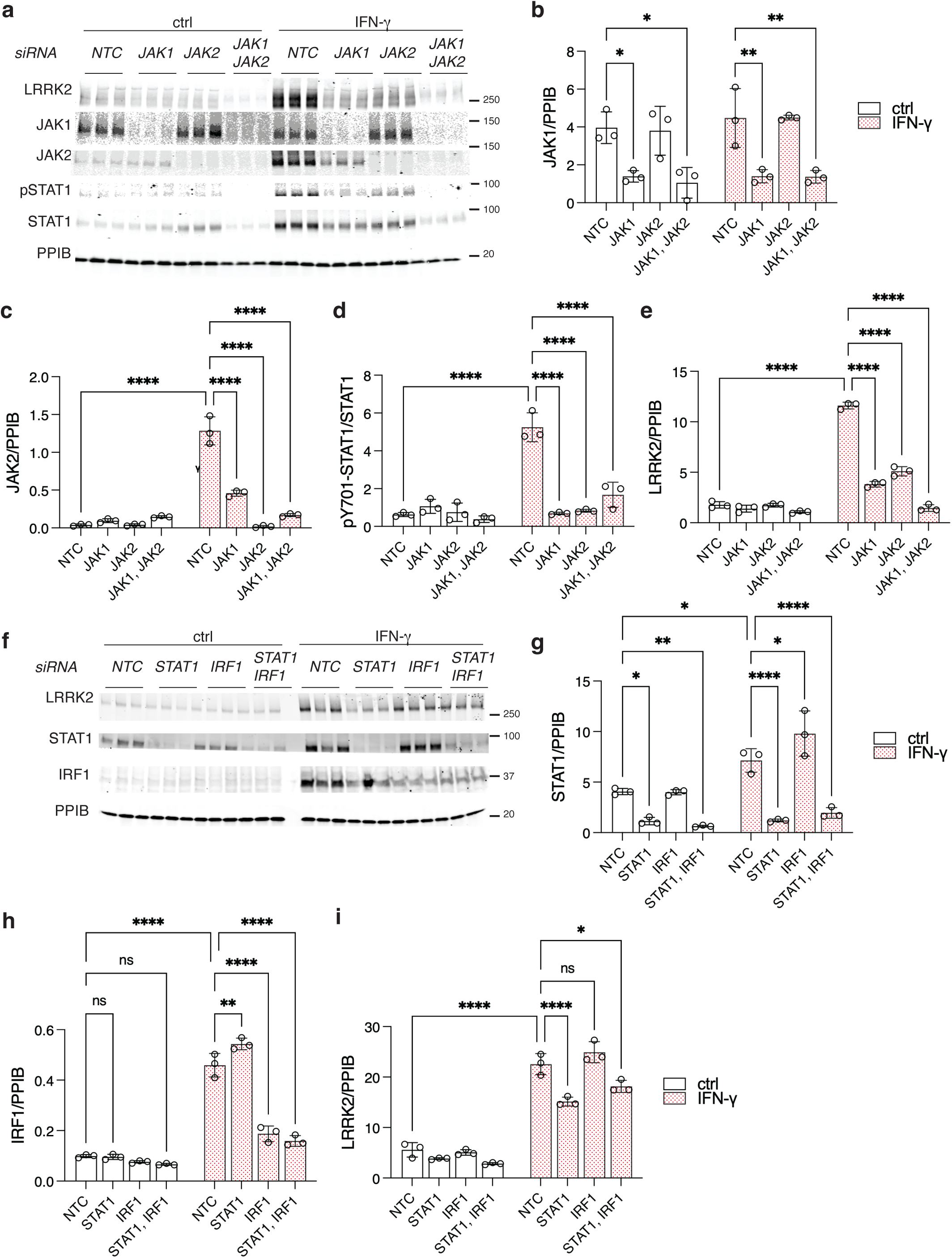
Validation of JAK/STAT signaling on LRRK2 expression after IFN-ɣ treatment. (a) Western blot analysis of JAK1 and JAK2 knockdown effects on LRRK2 induction. Knockdown of JAK1 and JAK2 resulted in a significant decrease in LRRK2 protein levels, with the most pronounced decrease observed upon combined knockdown of both kinases. Representative Western blot images are shown (a), along with quantification of JAK1 (b; F_1,16_ treatment = 1.227; F_3,16_ siRNA = 23.03), JAK2 (c; F_1,16_ treatment = 205.7; F_3,16_ siRNA = 94.46), phosphorylated STAT1 at Y701 (pY701 STAT1, d; F_1,16_ treatment = 65.03; F_3,16_ siRNA = 34.63), and LRRK2 (e; F_1,16_ treatment = 1001; F_3,16_ siRNA = 333.2) protein levels. Data are normalized to loading control (PPIB) or total protein (STAT1) and presented as mean ± SD with individual points showing replicate cultures. (f) Western blot analysis of IRF1 and STAT1 knockdown on LRRK2 induction. Knockdown of STAT1 but not IRF1 resulted in a significant reduction in LRRK2 protein levels Representative Western blot images are shown (f), along with quantification of STAT1 (g; F_1,16_ treatment = 45.22; F_3,16_ siRNA = 59.62), IRF1 (h; F_1,16_ treatment = 716.7; F_3,16_ siRNA = 120.8), and LRRK2 (i; F_1,16_ treatment = 911; F_3,16_ siRNA = 25.96) protein levels. Data are normalized to loading control (PPIB) and presented as mean ± SEM. \**p*<0.05; ** *p* <0.001; ***, *p*<0.0001 using two-way ANOVA to assess the effects of treatment and siRNA (values of F given above), followed by Tukey’s post-hoc test for multiple comparisons across all conditions. For all blots, each lane is a replicate culture and molecular weight markers are shown on the right. For all graphs, bars show mean values, error bars are standard deviation and individual points represent independent cultures.

**Figure S5.**
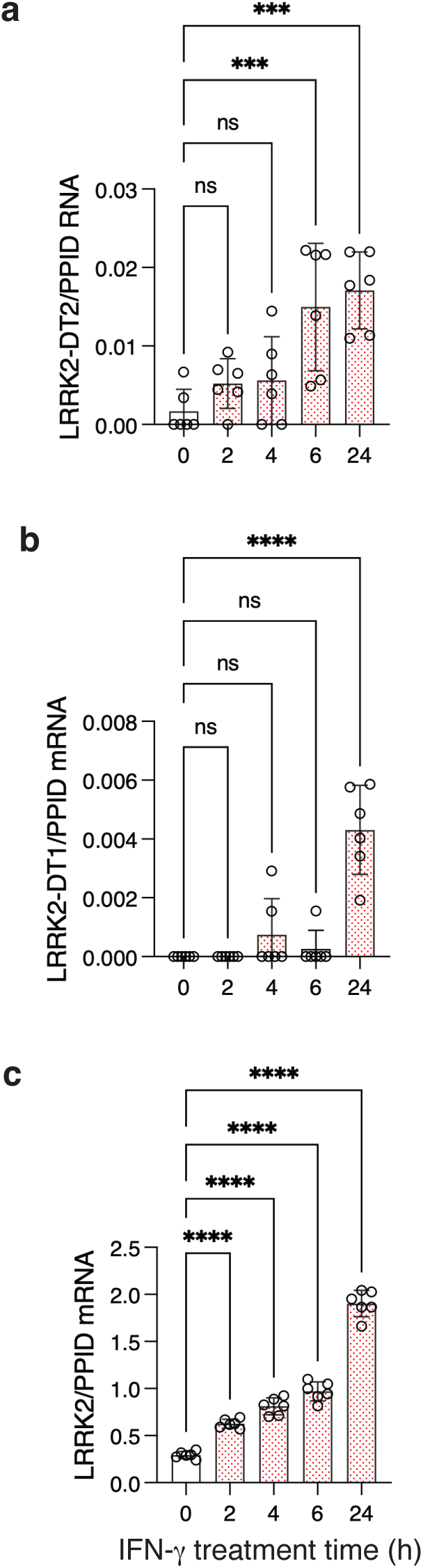
Time course of LRRK2 and LRRK2-DT mRNA induction following IFN-ɣ treatment. Quantitative PCR analysis of long non-coding RNAs LRRK2-DT1 and LRRK2-DT2, as well as LRRK2 mRNA, at indicated times after IFN-ɣ treatment. LRRK2-DT transcripts (a, b) showed a significant increase at 6h and 24 hours for LRRK2-DT2 and only at 24 hours for LRRK2-DT1, whereas LRRK2 mRNA levels (c) were significantly elevated as early as 2 hours post-treatment. ns, *p*>0.05; * *p*<0.05; ** *p* <0.001; ***, *p*<0.0001 using one-way ANOVA to compare treatments (F_4,25_ = 24.19 for LRRK2-DT1; F_4,25_ = 9.723 for LRRK2-DT2; F_4,25_ = 263.2 for LRRK2) followed by Dunnet’s post-hoc test for multiple comparisons compared to untreated cells. For all graphs, bars show mean value of expression relative to a reference mRNA PPID, error bars are standard deviation and individual points represent independent cultures.

**Figure S6.**
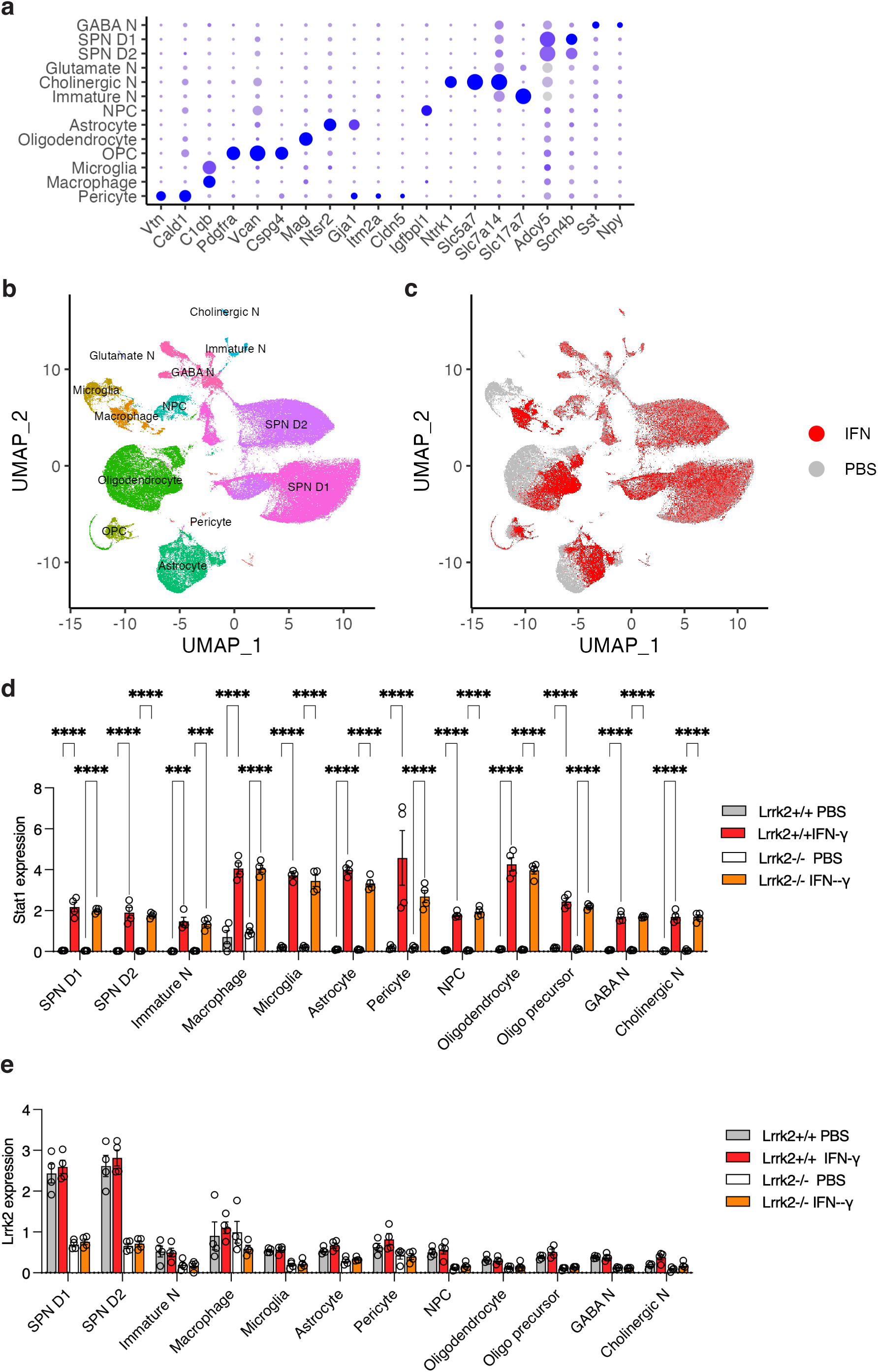
Interferon-γ does not significantly induce LRRK2 expression in the mouse striatum. (a) Dot plot analysis showing major cell subtypes in the striatum and associated GO term enrichment. Dot color intensity represents the average expression level of each gene within a cell type, while dot size indicates the proportion of cells expressing that gene. (b) UMAP visualization of major cell populations in the mouse striatum (150,360 cells). (c) UMAP comparison of striatal cells from PBS- and Ifn-ɣ injected mice. (e) Stat1 expression across major cell types in the striatum, comparing ± Ifn-ɣ treatment and LRRK2 wild-type (WT) versus knockout (KO) mice. (f) LRRK2 expression across major cell types in the striatum, comparing PBS and Ifn-ɣ treatment in LRRK2 WT versus KO mice.\**p*<0.05; ** *p* <0.001; ***, *p*<0.0001 by three-way ANOVA for Ifn-ɣ exposure by genotype and by cell-type (For Stat1 - F_10,132_ for cell type = 25.56; F_1,132_ for genotype = 4.483; F_1,132_ for IFN-ɣ exposure = 1360; For Lrrk2 - F_10,132_ for cell type = 97.49; F_1,132_ for genotype =319.2; F_1,132_ for IFN-ɣ exposure = 2.494; n=4 mice per genotype and treatment group) with Tukey’s *post-hoc* test for individual comparisons; for clarity, only comparisons between PBS and IFN-ɣ for each cell type that were significant (p<0.05) are shown.

**Figure S7.**
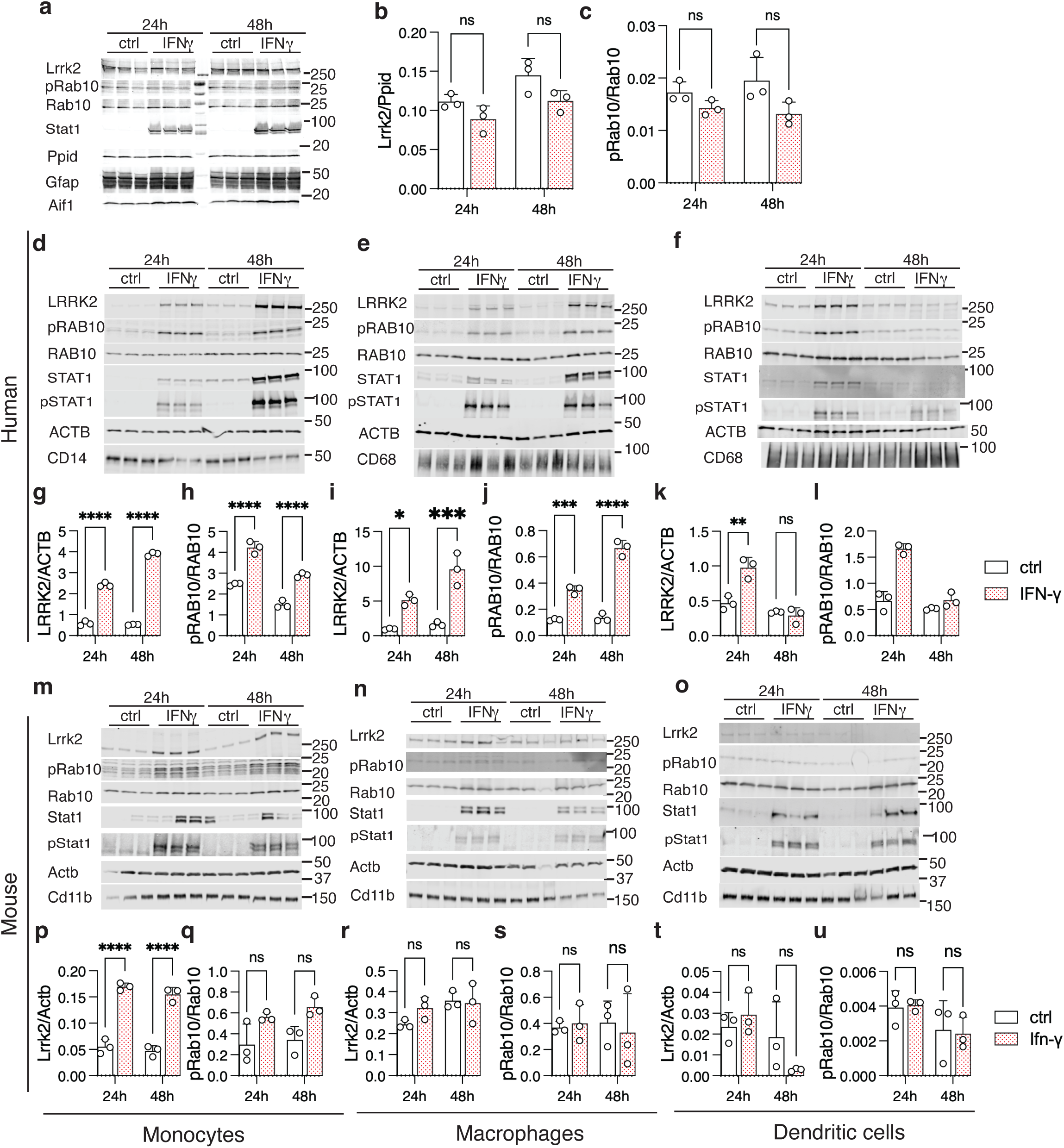
Interferon-γ induces weaker LRRK2 expression in murine than in human monocyte-lineage cells. (a) Mixed mouse glial cultures were blotted for untreated (ctrl) or exposed to mouse interferon-𝛾 for 24 or 48 hours and blotted for Lrrk2, pRab10, total Rab10, Stat1, pStat1 (Y701) the reference protein Ppib and makers for astrocytes and microglia (Gfap and Aif1). Quantification showed a modest decrease in Lrrk2 (b, F_1,8_ treatment = 9.14, *p*=0.016) and pRab10 (c, F_1,8_ treatment = 8.35, *p*=0.02) although no comparisons between treated and untreated cells at each time point were significant (Tukey’s post-hoc test all comparisons, all p>0.05, n=3 independent cultures per treatment and timepoint). Panels (d-f) show representative western blots for human monocytes (d), macrophages (e), and dendritic cells (DCs) (f) respectively, for LRRK2, pRAB10, RAB10, STAT1, ACTB, CD14, CD68, and CD11b protein levels after 24 and 48 hours of IFN-ɣ treatment in different donors from those in figure 6. Quantification of protein expression is shown in human monocytes (g, LRRK2; F_1,8_ treatment = 2476; h, pRAB10; F_1,8_ treatment = 217.2), macrophages (i, LRRK2; F_1,8_ treatment = 69.1; j, pRAB10; F_1,8_ treatment = 317.5) and dendritic cells (k, LRRK2; F_1,8_ treatment = 14.52; l, RAB10; F_1,8_ treatment = 66.3). Panels (m-u) show representative western blots for mouse monocytes (m), macrophages (n), and DCs (o), respectively in independent donor mice from those shown in figure 6. Quantification of protein expression is shown in mouse monocytes (p, Lrrk2; F_1,8_ treatment = 282.1; q, pRab10; F_1,8_ treatment = 15.0), macrophages (r, Lrrk2; F_1,8_ treatment = 0.97; s, pRab10; F_1,8_ treatment = 0.8423) and dendritic cells (t, Lrrk2; F_1,8_ treatment = 0.65; u, pRab10; F_1,8_ treatment = 0.00). In all graphs, ns p>0.05, \**p* < 0.05; **p < 0.001; \*\*\**p* < 0.0001 by two-way ANOVA assessing treatment and time point (values of F given above), followed by Tukey’s post-hoc test for pairwise comparisons. For clarity, only comparisons between control and interferon exposure are shown within each time point using n=3 replicate cultures per treatment and time point.

**Figure S8.**
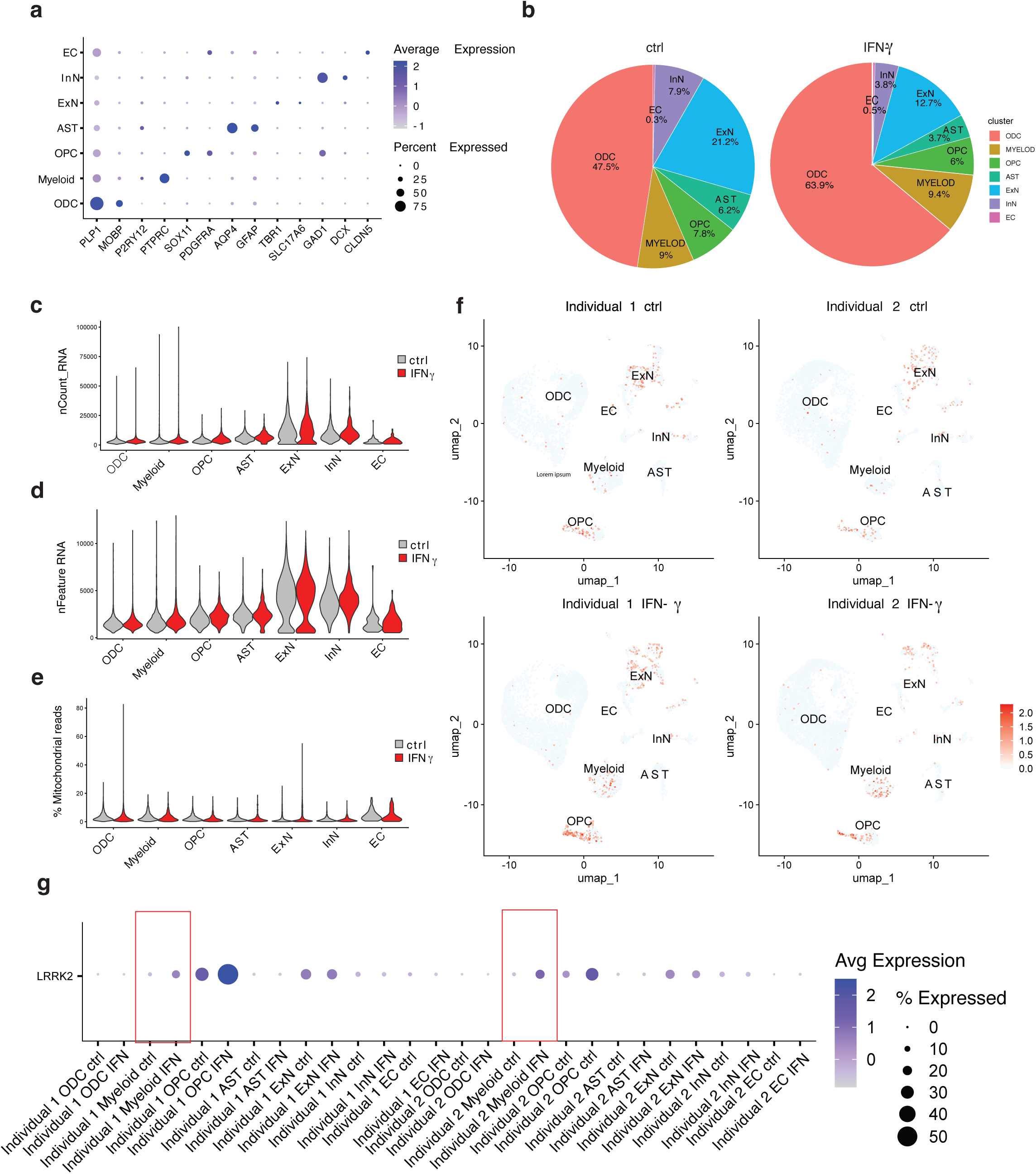
LRRK2 is induced by interferon-γ in human brain slices ex vivo. (a) Dot plot analysis showing major cell subtypes in the human brain sections and associated GO term enrichment. Dot color intensity represents the average expression level of each gene within a cell type, and dot size reflects the proportion of cells expressing that gene. (b) Pie chart illustrating the percentage of cells corresponding to major cell subtypes, showing conditions with and without IFN-ɣ treatment. (c–e) Quality control (QC) metrics by sample, including total RNA counts (nCount RNA, c), number of detected genes (nFeature RNA, d), and percentage of mitochondrial DNA reads (e). (f) Feature plot showing LRRK2 expression with and without IFN-ɣ treatment for each individual sample. The scale bar indicates relative expression levels. (g) Quantification of LRRK2 expression across major cell types in human brain sections for each individual, comparing without or with IFN-ɣ treatment. A total of 36,526 cells were used for analysis: 18,440 cells from control samples and 18,086 cells from IFN-γ–treated samples.

**Figure S9.**
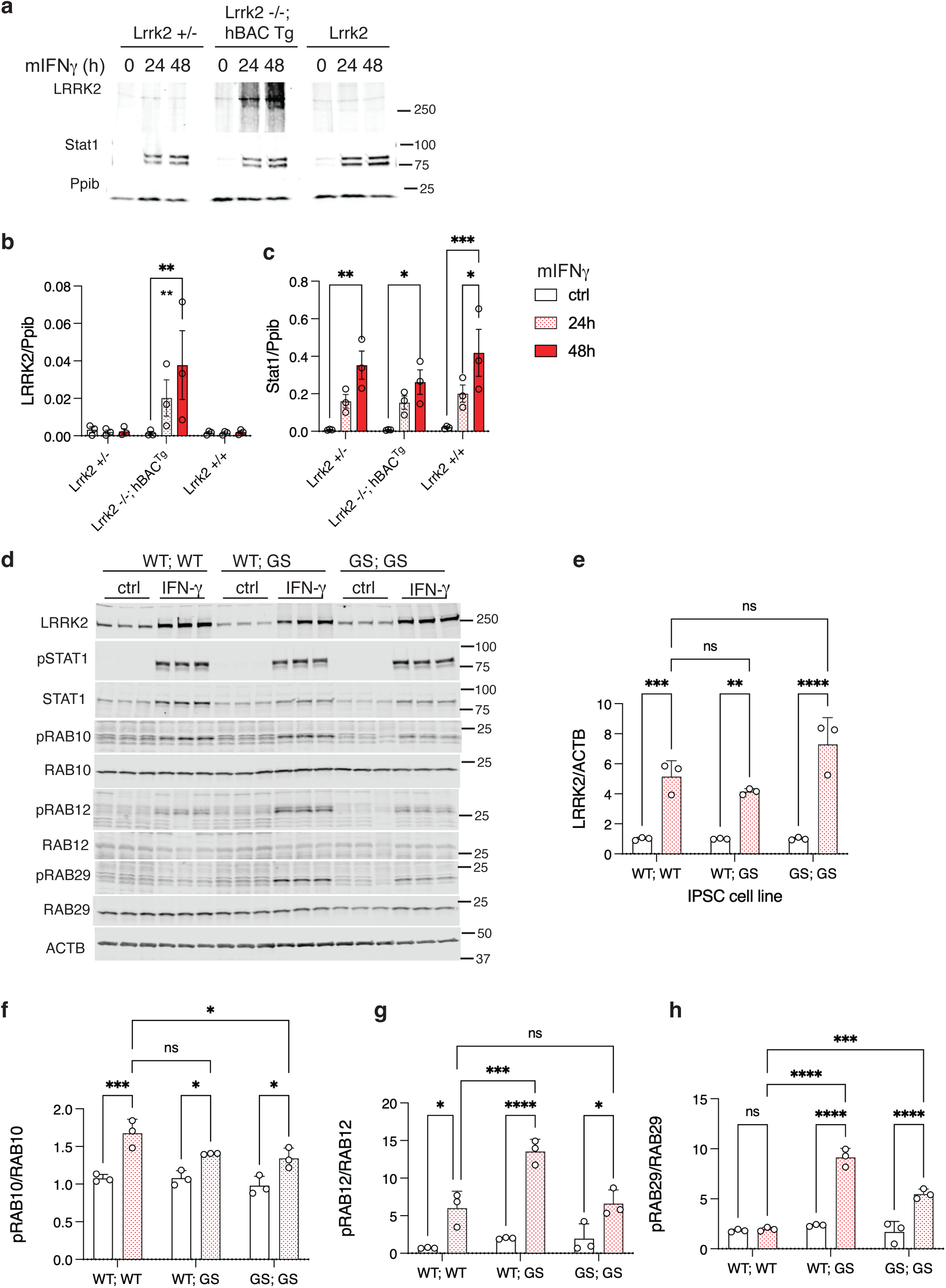
Combined effects of interferon-γ one human LRRK2 in mouse microglia and the LRRK2 p.G2019S mutation in human iMicroglia. (a-c) Relative induction of mouse and human Lrrk2/LRRK2 in primary mouse microglia. (a) Primary microglia were made from heterozygous mice (Lrrk2 +/-. i.e. one mouse allele), Lrrk2 knockout mice carrying a single copy of the human LRRK2 BAC (Lrrk2 -/-; hBAG Tg, ie one human allele) or wild type mice (Lrrk2 +/+) and blotted for LRRK2 (both human and mouse combined), Stat1 or Ppib (cyclophilin-b) as a loading control. Quantification shows strong induction of LRRK2 ony in microglia derived from mice carrying human LRRK2 but not other genotypes (b) which Stat1 was induced in all genotypes (c). For LRRK2 induction - F_2,18_ treatment = 2.2 ; F_2,18_ genotype = 6.5; F_4,18_ interaction = 2.3; For pStat1 – F_2,18_ treatment = 24.56; F_2,18_ genotype = 1.16; F_4,18_ interaction = 0.4344; with Tukey’s *post-hoc* test to compare all conditions. (d-h) LRRK2 induction by IFN-γ in human iMicroglia derived from wild-type, heterozygous (WT/GS), or homozygous (GS/GS) endogenous p.G2019S alleles (A18945 background). (d) Following protein extraction, immunoblots were performed for LRRK2, pSTAT1, STAT1, pRAB10, pRAB12, and pRAB29, along with appropriate loading controls. LRRK2 protein levels were increased by IFN-γ regardless of genotype (e). *, *p*<0.05; ** *p* <0.001; ***, *p*<0.0001 by two-way ANOVA for treatment by genotype (For LRRK2 induction – F_1,12_ treatment =.127.7 ; F_2,12_ genotype = 5.36; F_2,12_ interaction =5.36). pRAB10 induction was similar across all lines (f), whereas induction of pRAB12 (g) and pRAB29 (h) was higher in G2019S mutant lines. For pRAB10 - F_1,12_ treatment = 61.9 ; F_2,12_ genotype = 5.3; F_2,12_ interaction = 2.5; For pRAB12 - F_1,12_ treatment = 92.95; F_2,12_ genotype = 13.06; F_2,12_ interaction = 8.78; For pRAB29 – F_1,12_ treatment = 154; F_2,12_ genotype =60.4 ; F_2,12_ interaction = 44.68, with Tukey’s *post-hoc* test to compare all conditions. For all blots, each lane is a replicate culture and molecular weight markers are shown on the right. For all graphs, bars show mean values, error bars are standard deviation, and individual points represent independent cultures. N = 3 independent differentiations; representative of 3 independent experiments.

**Figure S10.**
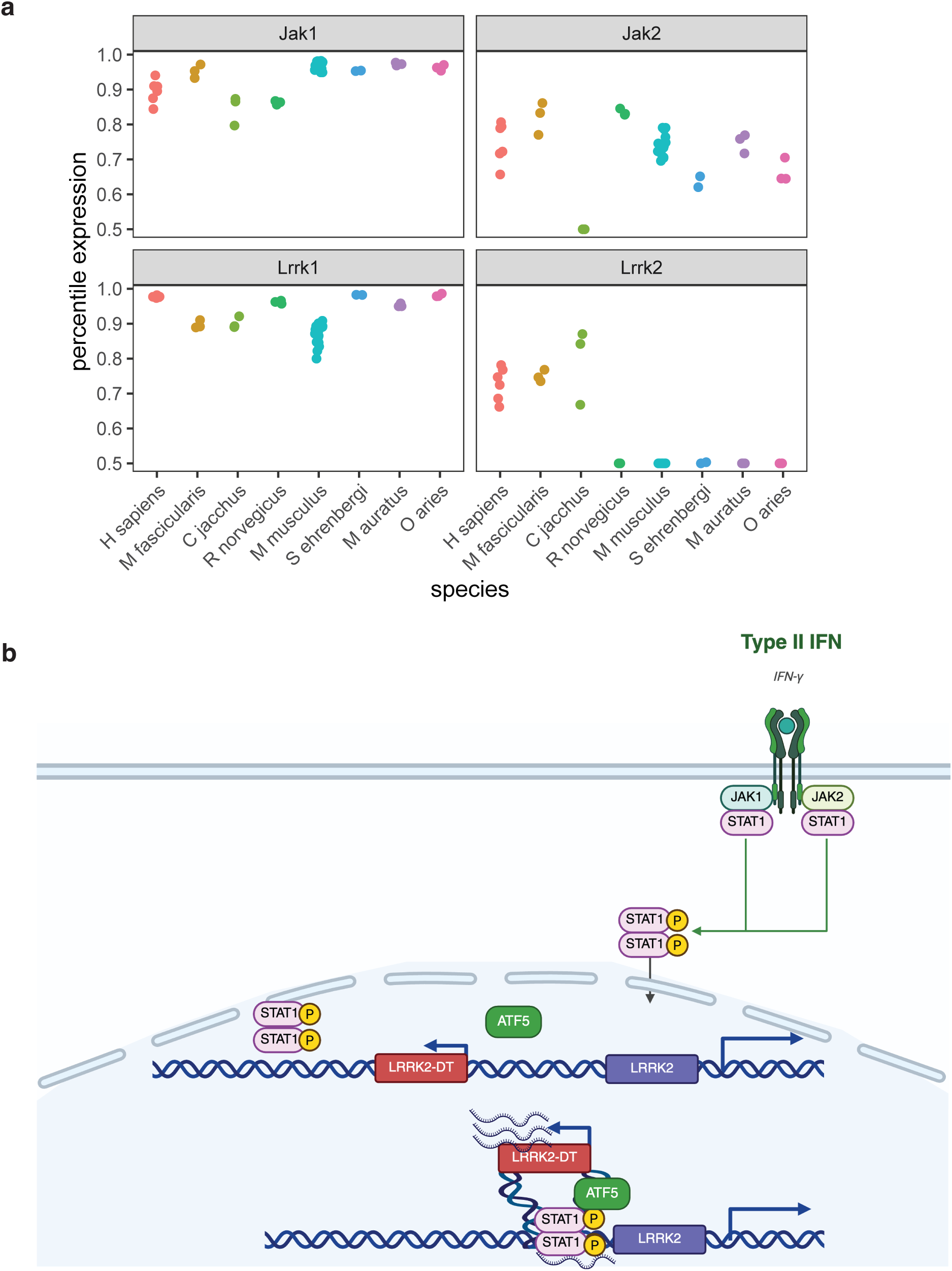
Human specific regulatory mechanisms for LRRK2 expression by interferon-γ. (a) Reanalysis of published data in isolated microglia (*35*) across species on the horizontal axis compared to percentile expression of the indicated genes on the vertical axis. While Lrrk2 expression is higher in humans and non-human primates compared to other mammals examined, Jak1, Jak2 and Lrrk1 are relatively consistently expressed across species. Each dot represents a single sample. (b) Schematic representation of the human specific regulation of LRRK2 by interferon-ɣ signaling.

## Notes

### Competing Interest Statement

The authors have declared no competing interest.

